# Astrocyte-specific secretome profiling reveals its correlation with neurological disorders

**DOI:** 10.1101/2024.11.21.624658

**Authors:** Jie Liu, Jing Gao, Lin Guo, Guoming Ma, Mingfeng Guan, Congcong Xia, Jia He, Yufang Yang, Yi Wu, Jun Xu, Liulin Xiong, Chang-Yin Yu, Gang Pei, Jian Zhao, Jianchen He, Yaoyang Zhang, Wenyuan Wang

**Affiliations:** Interdisciplinary Research Center on Biology and Chemistry, Shanghai Institute of Organic Chemistry, Chinese Academy of Science, Shanghai, 200032, China; The Public Experiment Platform, School of Basic Medical Sciences, Shanghai University of Traditional Chinese Medicine, 1200 Cailun Road, Shanghai 201203, China; Department of Rehabilitation Medicine, Huashan Hospital, Fudan University, Shanghai 200040, China; East Hospital, Tongji University School of Medicine, Shanghai, 200120, China; Department of Anesthesiology, Affiliated Hospital of Zunyi Medical University, Zunyi, 563006, China; State key laboratory of cell biology, CAS center for excellence in molecular cell science, Shanghai Institute of biochemistry and cell biology, Chinese Academy of Science, Shanghai, 200032, China; Shanghai Institute for Advanced Immunochemical Studies, ShanghaiTech University, Shanghai, 201210, China; Department of Diagnostics of Traditional Chinese Medicine, School of Basic Medical Sciences, Shanghai University of Traditional Chinese Medicine, 1200 Cailun Road, Shanghai, 201203, China; University of Chinese Academy of Sciences, Beijing, 100049, China; Animal Center of Zoology, Institute of Neuroscience, Kunming medical University, Kunming, 650500, China

**Keywords:** Astrocyte, Secretome, Click chemistry, Neurological disorders, non-cell autonomous toxicity

## Abstract

Secreted proteins mediate intercellular communication throughout the lives of multicellular organisms. However, due to the lack of new technology for secreted protein capturing, the progress of “secretomics’’ lags behind. Here, we report a two-step secretome enrichment method (tsSEM) combining unnatural amino acid labeling and click chemistry-based biorthogonal reaction, which enables *in vitro* secretome profiling in the presence of serum. Using this novel method, we systematically investigated the secretome of human iPSCs-derived astrocytes (iAst) in different disease models and identified a panel of astrocyte-secreted proteins that are responsible for its non-cell autonomous toxicity under disease conditions. Furthermore, we validated two astrocytes-derived novel neurotrophic proteins, FAM3C and KITLG, which we identified from disease models, and found that they could boost neurite outgrowth, protect neurons, and promote neural progenitor proliferation. Our study highlights the utility of secretome profiling of iAst and demonstrates its applications in disease study and target identification and validation in drug development.

## Introduction

The secretome constitutes an important class of proteins and organic molecules secreted by a cell, tissue, or organism and circulating throughout the body, modulating the extracellular environment, cell-cell communication, signal transduction, and many cellular functions in a paracrine or autocrine fashion. Secretory proteins have been widely implicated in various human diseases, including cancer metastasis, metabolic diseases, and neurodegenerative diseases^1^.

Astrocytes, the specialized glial cells in the brain, play multiple roles in maintain brain homeostasis, such as providing energy substrates for neurons, regulating neurotransmitter levels, modulation blood-brain barrier stability, *et al*. Under disease conditions, such as injury, infection, or neuroinflammation, astrocytes could transform from “neurotrophic” to “neurotoxic” states, release harmful substances such as inflammatory cytokines, chemokines, and neurotoxins, which can worsen inflammation and affect neuronal functions, ultimately leading to neuronal death^2,3^. As such, the neurotoxic substances secreted by astrocytes have been associated with the development of various neurological diseases^2^. Therefore, analysis of the astrocyte secretome would provide essential information to promote our understanding of the neurological diseases and open new avenues of novel diagnosis and therapeutic intervention.

Despite the importance of the astrocyte secretome, the research progress on it is strongly hampered by technical limitations. Due to the low abundance of secreted proteins against a background of highly abundant serum proteins, secretome profiling is conventionally collected under a serum-free condition^4–7^. However, such culture conditions usually render the cells into a disease-like state, especially for sensitive astrocytes, which can profoundly affect the proteomic profiling of the cells^8^. Additionally, our current understanding of astrocyte physiology is almost based on murine models, whose functional and molecular characteristics differs a lot to human astrocytes^9^. Fortunately, the derivation of human induced pluripotent stem cells (iPSCs) over a decade ago has overcome many limitations of animal models in neurogenerative disease research^10^. The iPSCs can be readily differentiated into many subtypes of functional brain cells for precise disease modeling, drug screening, and mechanism studies^11^. Astrocytes that are differentiated from iPSCs/ESC can be a valuable tool for investigating the mechanisms underlying neurological disorders. In this regard, analyzing the secretome of iPSCs-derived astrocytes (iAst) can provide crucial insights into the intercellular communication in the nervous system and associated disorders.

In the current study, we report an optimized method to precisely capture secreted proteins by combining azidohomoalanine (AHA) protein labeling and click chemistry. The labeled secreted proteome was processed by a tandem enrichment procedure and then subjected to a shotgun proteomic analysis. Using this optimized method, we systematically profiled the secretome of murine astrocytes (mAst), healthy individual, and AD patient iPSCs-derived iAst, respectively, and identified a panel of secreted proteins that responsible for the non-cell autonomous toxicity of AD patient iPSCs-derived iAst. We further validated two novel astrocyte-secreted neurotrophic proteins, FAM3C, which could protect neurons from death and promote the complexity of neurite branches; and KITLG, which could enhance neural progenitor cell proliferation and reduce neuronal death.

Collectively, our study not only provides rich resources for studying the cellular function of astrocytes for the research community but also demonstrates the power of the secretome profiling method in therapeutic target identification, validation, and mechanism studies. The data can open new avenues for novel diagnosis and therapeutic intervention for neurodegenerative diseases.

## Results

### Establishment of a two-step peptide level enrichment procedure

To profile astrocyte secretome in serum-containing media, we applied an alternative strategy by pulse labeling astrocytes with azidohomoalanine (AHA), an azide-bearing analog of methionine (Met) (Fig.S1A), which allowed the selective labeling of newly synthesized and secreted protein in the conditioned media, without affecting cell morphology, proliferation rate, and viability (Fig.S1B-D). The protein expression pattern was similar between the astrocytes grown in AHA replacement and regular medium (Fig.S1E, F). Meanwhile, 21.38% of Met-containing peptides were labeled with AHA after 24h of AHA incubation (Fig.S1G).

We first tested a published method^12^ for AHA-labeled protein isolation that allows the covalent capture of AHA-labeled proteins (Fig.S2A). Our pilot experiments revealed that the method did not meet our expectations in terms of specificity. With this method, about 20% (114 of 612) of the detected proteins were due to unspecific binding (Fig.S2B, C). To achieve specific AHA-labeled protein enrichment, we linked AHA-labeled proteins to alkyne-(PEG)4-biotin through click chemistry instead of covalently linking it to alkyne-activated resin (Fig.1A, upper left), and then isolated these proteins through the biotin-avidin system. We further developed a two-step enrichment procedure to improve the specificity and efficiency (Fig.1A, detailed in *Methods*). The advantage of this method is that it can recover AHA modified peptides by acid elution. This allows us to discriminate whether the protein is a specifically enriched secretory protein with the information of AHA modification in subsequent data analysis.

**Fig.1.**
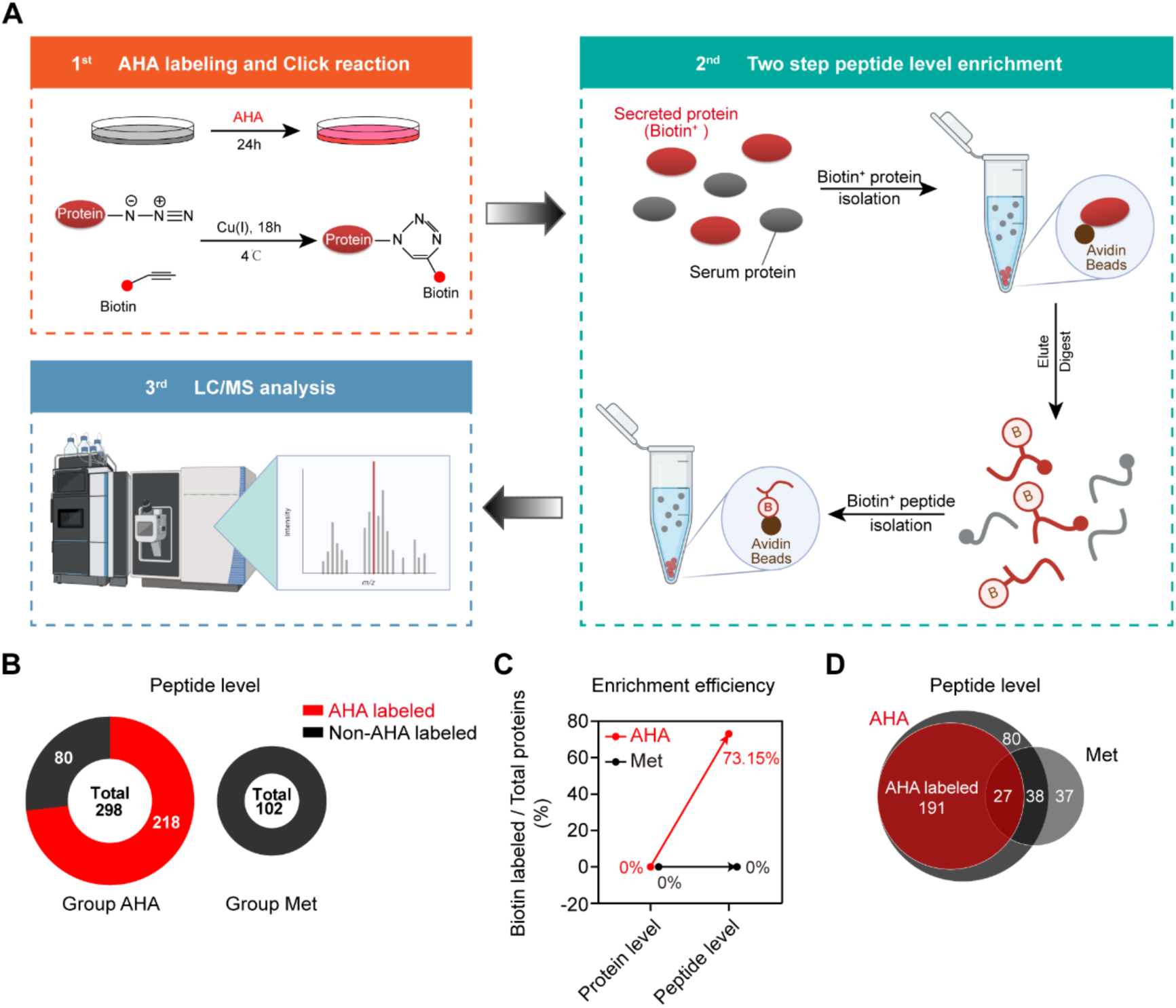
Establishment of a two-step peptide level enrichment procedure. A, Schema of a two-step peptide enrichment procedure. In brief, after AHA labeling and click reaction, biotinylated proteins were pulled down using Avidin resin. Instead of directly on-bead digestion, we next eluted these resin-binding proteins, digested them into peptides, and conducted the second step of peptide level enrichment. These resin-binding peptides were eluted and processed for LC/MS analysis. B Numbers of proteins profiled by two-step peptide enrichment. C, Enrichment efficiency of two different methods. Enrichment efficiency is calculated as AHA-protein/Total protein. D, Comparison of the profiled proteins between the AHA and Met group.

We applied this method for the secretome profiling. In the AHA-replacement culture group, 298 proteins were purified (Fig.1B), 73.15% of which were AHA labeled, much higher than the previous method (0%, Fig.1C). In the Met group, we detected 102 proteins, 65 of which were shared with the AHA group (Fig.1D). In the previous method, the proteins shared between Met and AHA groups had to be discarded as non-specific binding proteins. With the advantage of our method, we could clearly distinguish secreted proteins from unspecific binding proteins and preserve peptide information as much as possible.

Taken together, we have developed a highly specific and efficient two-step secretome enrichment method (tsSEM) that combines unnatural amino acids (AHA) incorporation and click reaction. This allows us to identify secreted proteins in serum-containing conditions specifically.

### Secretome profiling of primary mice astrocytes

We first applied this method for primary mice astrocytes (mAst) secretome profiling. About 300 biotin-labeled peptides were detected (Fig.2A). The enrichment efficiency was above 80% (Fig.2B). These peptides correspond to a total of 192 proteins (Table S1). Gene Ontology analysis indicates that the enriched proteins were mainly distributed in the extracellular compartment (Fig.2C). Subcellular localization annotation revealed that more than 85% of the enriched proteins were secreted proteins (Fig.2D). This ratio was much higher than that of traditional serum-free culture conditions (less than 50%)^13^. Furthermore, according to the DAVID knowledgebase, 49% of the identified proteins are highly expressed in the brain (94 out of 192, Fig.2E), which coincides with the brain origin of astrocytes.

**Fig.2.**
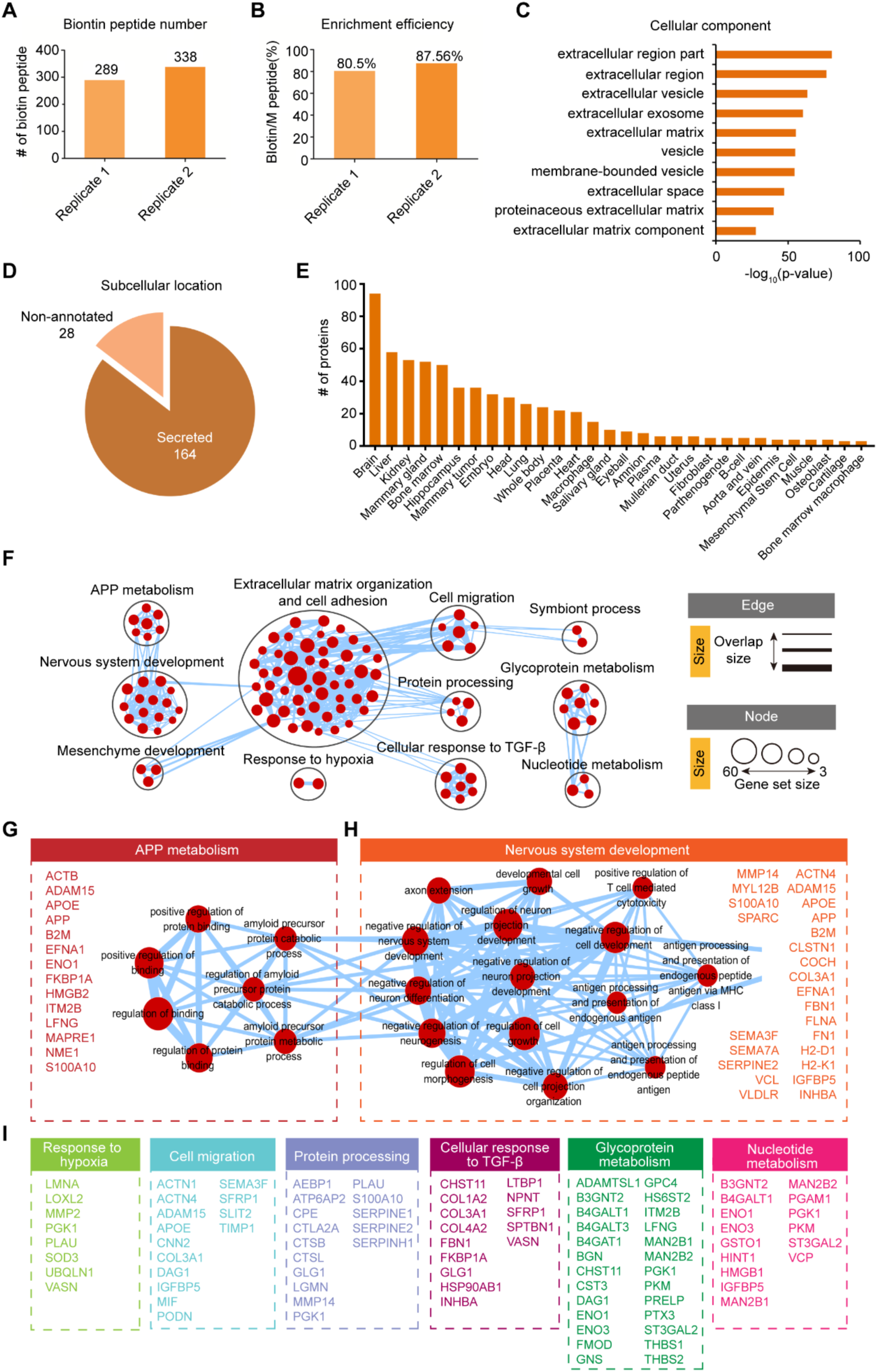
Profiling of mice primary astrocytes secretome. A, B, mAst secretome was analyzed in duplicate by LC-MS/MS; presented are the enriched biotin-labeled peptide number (A) and enrichment efficiency (B) of each replicate. N=2 biological replicates. Given the large number of cells required for each replicate, about 2.5×10^7^ cells which collected from ten 15-cm dish, we have only designed 2 sets of biological replicates for each profiling. C, Cellular component distribution of the profiled 192 mAst secretome proteins was analyzed by the DAVID database; the top 10 significant terms were all related to extracellular location. D, Statistical analysis of the subcellular location of mAst secretome, proteins annotated as secreted, extracellular space, and cell membrane in the Uniprot database were all regarded as secreted. E, Tissue distribution of mAst secretome according to the DAVID database. F, Pathway enrichment analysis of 192 mAst secretome proteins, the dataset was analyzed by g: Profiler and further processed with Cytoscape. G-H, The detailed pathways and proteins enriched in APP metabolism(G) and nervous system development(H) clusters. I, Proteins enriched in response to hypoxia, cell migration, protein processing, cellular response to TGF-β, glycoprotein metabolism and nucleotide metabolism clusters.

Next, we compared the top 500 astrocytes uniquely expressed genes reported by Zhang *et al* ^14^ with our mAst secretome and found that 10 out of 192 secreted proteins were highly expressed in astrocytes at the transcriptional level (Fig.S3). Among these proteins, pentraxin-related protein (PTX3) and latent-transforming growth factor beta-binding protein1 (LTBP1) are involved in immune response and regulate inflammation in response to infectious conditions^15^. PTX3 is also critically involved in synaptic function by inducing AMPA receptor clustering^16^. Astrocytes secreted glypican-4 (GPC4) can regulate the release of neuronal pentraxin1 from axons to induce functional synapse formation^17^. Thrombospondin-1 (THBS1) is able to promotes synaptogenesis and modulates synapse and spine defects in the fragile X mouse model^18,19^. These studies collectively demonstrated the critical role of astrocyte-secreted proteins in synaptogenesis and neural plasticity modulation. Further investigation of these proteins will expand our understanding of the function of astrocytes in synapse development and diseases.

The function of astrocytes has evolved rapidly in the past two decades from “brain glue” to a scaffold necessary for neuronal distribution and interaction, and more recently, play essential roles in neurodevelopment, synapse formation, intercellular communication, and neurological diseases ^20,21^. Indeed, functional annotation analysis indicates that the most enriched molecular pathway is extracellular matrix organization and cell adhesion (Fig.2F). Biological pathways associated with nervous system development, including neuronal differentiation, axon extension, cell morphogenesis, and projection development were also enriched (Fig.2F). Meanwhile, the secreted proteins participated in neurological diseases related pathways, such as APP metabolism, cellular response to TGF-β and hypoxia, and amyloid-β processing. For the APP metabolism pathway, 14 secreted proteins were clustered, including apolipoprotein E (APOE), amyloid-beta precursor protein (APP), integral membrane protein 2B (ITM2B), high mobility group protein B2 (HMGB2), and *et al*. APOE is mainly expressed and secreted by astrocytes in CNS, and the ε4 allele of the *ApoE* is the most significant genetic risk factor for late-onset sporadic AD ^22^. ITM2B has been shown to interact with amyloid precursor protein (APP) and regulate amyloid-β (Aβ) production^23^. Interestingly, some proteins are involved in AD pathology, but how they regulate APP metabolism has yet to be experimentally validated ^24^. For example, nucleoside diphosphate kinase A (NME1)^25^ has been found exclusively contained in post-mortem human brains affected by AD but not in non-AD extracts.

We further analyzed the protein-protein interaction (PPI) of the secreted proteins enriched in each cluster (Fig.S4). The result shows that these proteins form a very complex interaction network, and the most connected proteins are all well recognized in regulating the corresponding cellular pathways.

Taken together, we successfully profiled secretome from primary cultured mouse astrocytes, and the percentage of secreted proteins in our dataset is higher compared to the previous studies^4,6,13,26^. Pathway enrichment and PPI analysis revealed that these proteins are major effectors of the biological functions of astrocytes.

### Secretome profiling of human iPSC-induced astrocytes

To the best of our knowledge, there are still currently no comprehensive profiling of human astrocytes secretome. We thus investigated this issue using iPSC-derived human astrocytes with our tsSEM method. We optimized the protocol for astrocyte differentiation based on the method reported by Krencik R *et al.*^27^, which allowed us to obtain highly purify human iPSC-derived astrocytes (iAst) for secretome profiling (Fig.3A, B). We collectively identified 293 proteins (Fig.3C, Table S2), of which 88% were annotated as secreted proteins (Fig.3D-E). Based on the DAVID Knowledgebase, 143 out of 293 are expressed at a high level in the human brain (Fig.4F).

**Fig.3.**
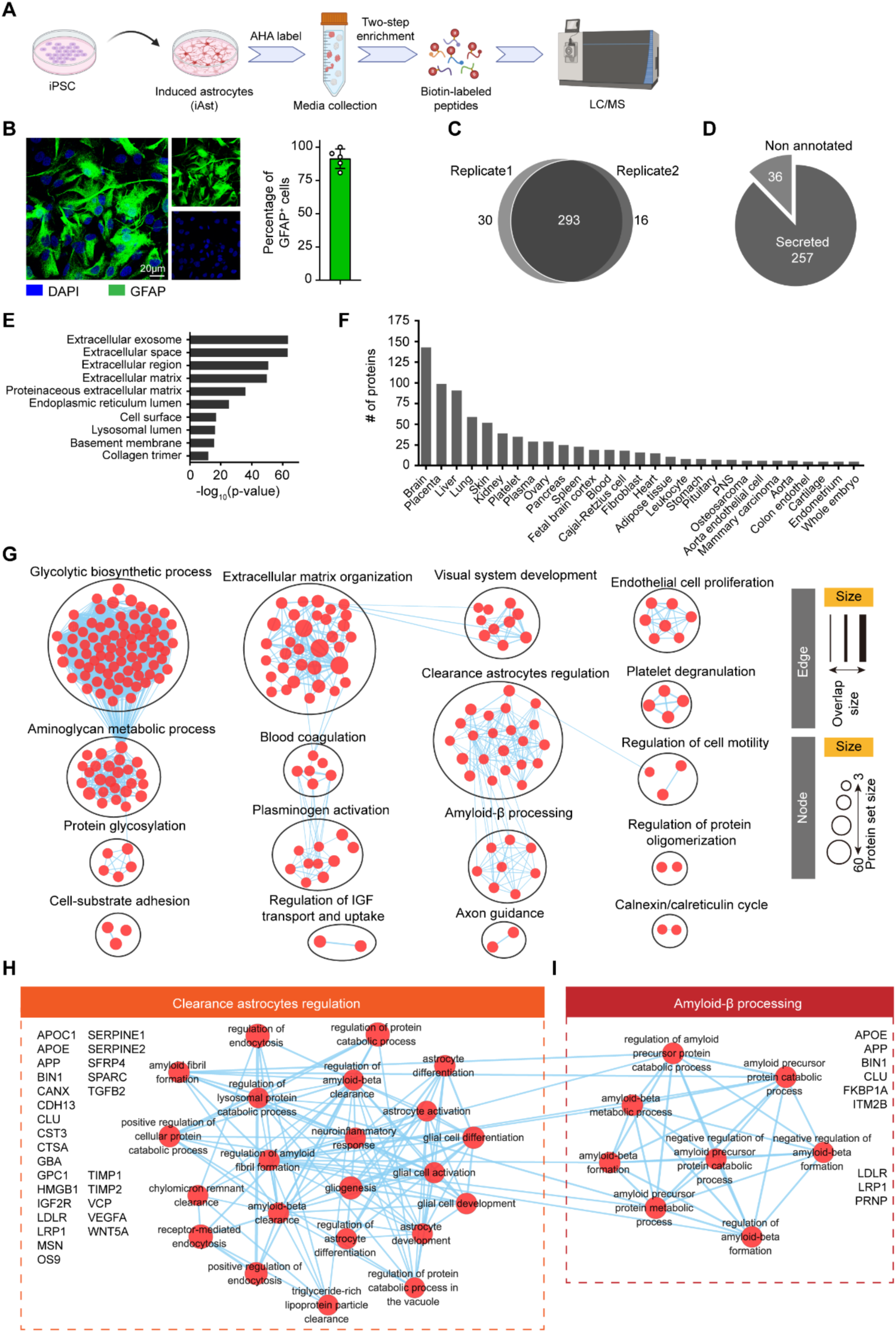
Profiling of human-induced astrocytes secretome. A, Schema of human-induced astrocytes secretome profiling. B, Representative images of induced astrocytes from a healthy human ESC line, iAst were labeled with GFAP (green) antibody. C, iAst secretome was analyzed in duplicate by LC-MS/MS, and more than 90% of the qualified proteins were found in both replicates. N=2 biological replicates. D, Statistical analysis of the subcellular location of secretome proteins, proteins were annotated as secreted, extracellular space and cell membrane in Uniprot database were all regarded as secreted in our statistics. E, Cellular component distribution of the overlap 293 secretome proteins was analyzed by DAVID database, almost all the top 10 enriched terms were related to extracellular location. F, Tissue distribution of iAst secretome according to the DAVID database. G, Pathway enrichment analysis of profiled secretome proteins, the dataset was analyzed by g: Profiler and further processed with Cytoscape. H,I, The detailed pathways and proteins enriched in Amyloid-β processing(H) and clearance astrocytes regulation(I) clusters.

**Fig.4.**
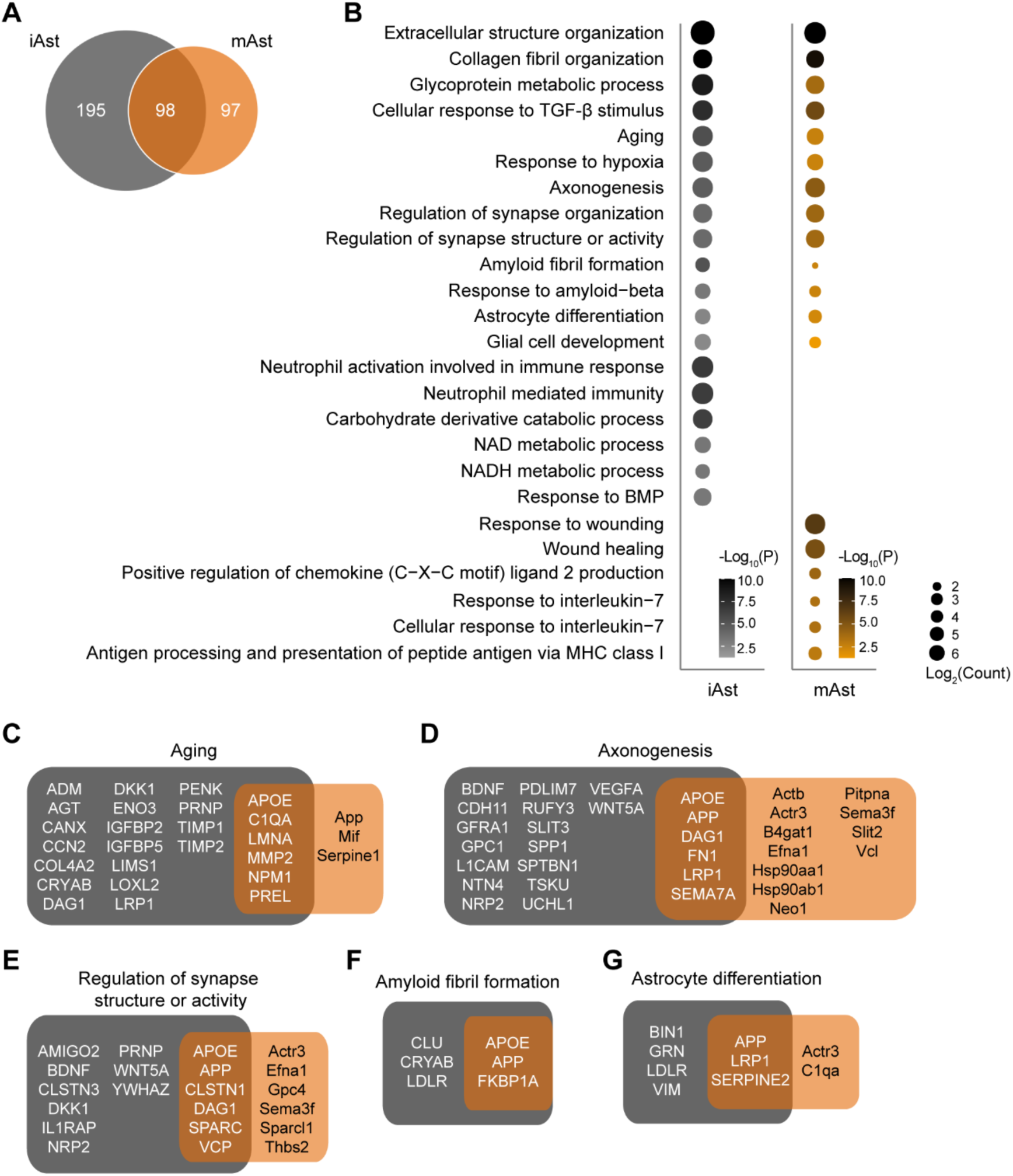
Comparison between iAst and mAst secretome. A, Comparison of the iAst and mAst secretome. N=2 biological replicates for each group. B, Gene ontology analysis of two sets of the secretome. The biological processes shared by the two groups, and the most significant enriched by each are shown here. C-G, Specific proteins enriched in the representative shared biological processes.

We compared the iAst secretome with the top 500 astrocyte-expressed genes (Fig.S5). We found that 15 out of 293 secreted proteins were also transcriptionally highly expressed in astrocytes. Additionally, among these 15 highly expressed proteins, four were also on the top secreted protein list, including IGFBP1 (regulates glial proliferation and survival) ^28^, MMP14 (a member of matrix metalloproteinase family implicated in neurodegenerative and neuro-inflammatory pathologies)^29^, LOXL1 (a lysyl oxidase essential for the formation of crosslinks between collagen and elastin, and implicated in exfoliation glaucoma and the progression of glioma)^30^ and LTBP1 (a key regulator of TGF-β signaling involved in astrocyte scar formation and astrocytes maturation)^31^. VEGFA and TGFB2 have already been reported to be secreted by astrocytes and participate in brain-blood-barrier (BBB) regulation^32^. BDNF and NOTCH3 are also well-known regulators of nervous system development^33,34^.

Pathway enrichment analysis indicates that the most significantly enriched cellular pathway is related to glycan biosynthesis and metabolism, including the glycolytic biosynthetic process, aminoglycan metabolic process, and protein glycosylation, suggesting the critical role of astrocytes in glycan homeostasis regulation (Fig.3G). Consistent with the mAst, iAst-secreted proteins are enriched in the extracellular matrix organization, development, and protein processing processes. Interestingly, molecular functions connected with neurodegenerative disorders, such as protein clearance and Aβ processing, are also enriched in iAst secretome. A closer look at these two clusters revealed that the secreted proteins of iAst mediate many critical subcellular functions closely related to protein homeostasis in neurological diseases, such as regulation of Aβ formation and clearance, APP catabolic process, neuro-inflammation response, gliogenesis, regulation of endocytosis, and lysosomal protein catabolic process (Fig.3H, I).

We conducted a PPI analysis to identify the hub proteins that may have a more significant functional role. As depicted in Fig.S6, the results have highlighted several well-known modulators of the enriched biological process. These include APP, APOE, and LDLR, which are associated with protein clearance and Aβ processing. IGFBP3, VCAN, CST3, and PENK, and multiple ADAMTS family members are involved in the glycolytic biosynthetic process and IGF transport/uptake. Additionally, MMP1, MMP3, and APOE are associated with protein oligomerization.

We then compared the secretome between the iAst and mAst. In general, we identified more secreted proteins from iAst than mAst (293 versus 192) (Fig.4A). The proteins shared by the two groups are mainly enriched in the fundamental biological processes of astrocytes, such as extracellular matrix organization, axon guidance and regulation of synapse organization (Fig.4B). iAst-specific secreted proteins were also enriched in some conventional metabolic processes. In comparison, the mAst-specific secreted proteins were enriched in inflammatory response processes. We reasoned that mouse astrocytes were isolated from postnatal mouse brains, the *in vivo* developmental stage of mAst may render the cells more sensitive to extracellular stimulation. We also established the specific proteins enriched in several biological processes. The number of proteins involved in these processes is higher in the iAst group, which to a certain extent indicates that the function of human astrocytes is more complicated than that of mice (Fig.4C-G), and our secretome dataset can well reflect this difference.

### Secretome profiling of iAst under TNF-α stimulation

Astrocytes are the critical regulators of neuroinflammation. So, we profiled the secretome of iAst treated with TNF-α, one of the pro-inflammatory cytokines involved in the pathogenesis of various neurological diseases. We identified a total of 249 proteins after TNF-α treatment (Fig.S7A, Table S3), and most of them are annotated as secreted proteins (Fig.S7B, C); 120 out of 249 are expressed in the highest level in the brain according to the DAVID knowledgebase (Fig.S7D). 43 proteins were uniquely identified in TNF-α treated iAst (Fig.5A), which mainly participate in inflammation related biological processed, such as antigen processing and presentation, pathogenic E. coli infection and regulation of apoptotic process (Fig.5B), indicating that iAst secreted proteins may be the mediators of these pathological effects.

**Fig.5.**
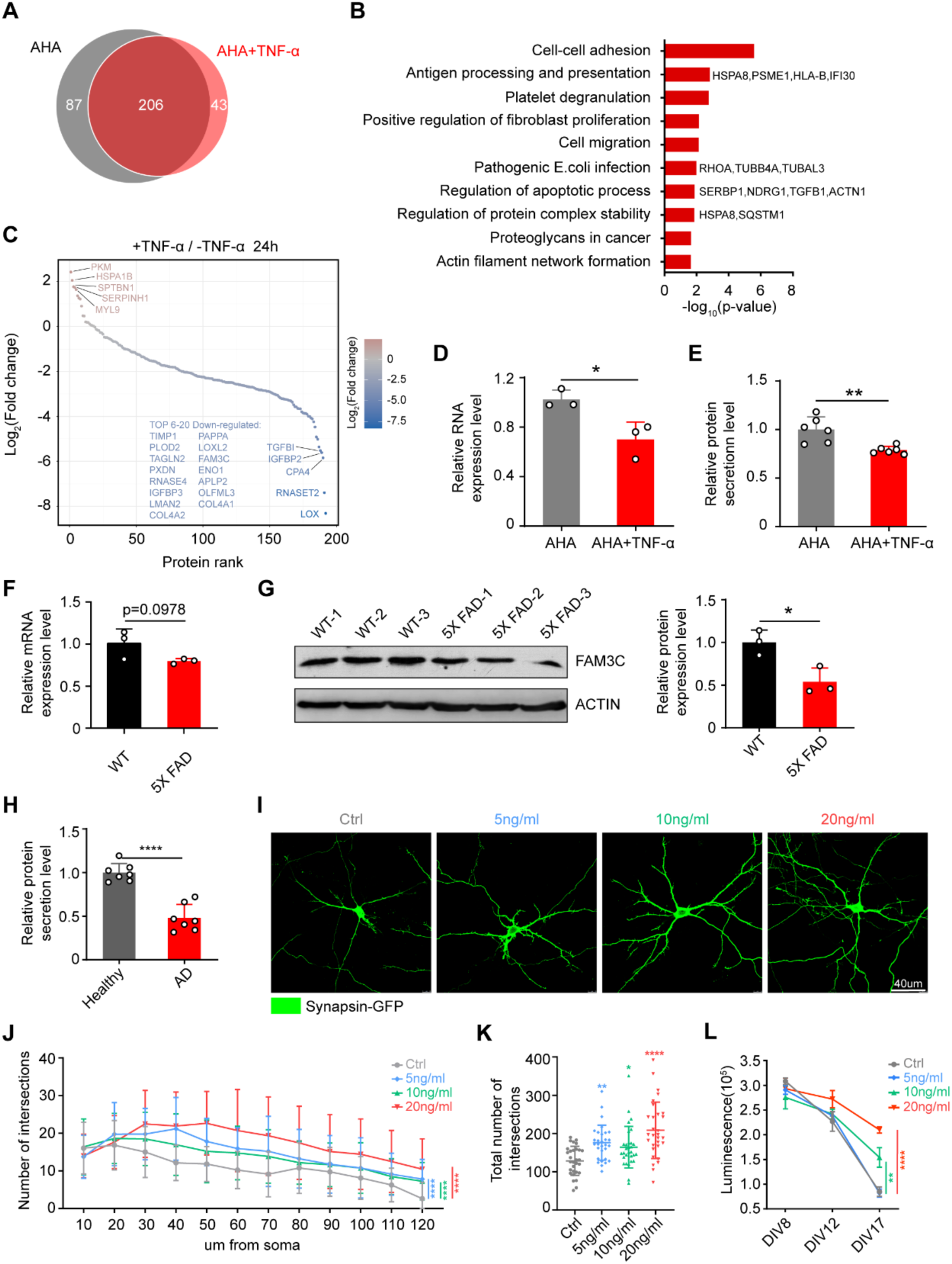
Impactions of TNF-α treatment on iAst secretome. A, The secretome of TNF-α (30ng/mL) treated iAst consists of 43 unique proteins and 206 overlapped proteins compared to vehicle-untreated iAst secretome. N=2 biological replicates for each group. B, According to the DAVID database, Top 10 enriched biological processes of the TNF-α treated astrocytes uniquely secreted proteins. C, Relative quantitative analysis of the 206 overlapped secreted proteins. D, Relative iAst FAM3C RNA expression level change after TNF-α treatment. Data are presented as mean ± SD, n=3. Two-tailed unpaired t-test, *p<0.05. E, Relative iAst FAM3C secretion level change after TNF-α treatment. Data are presented as mean ± SD, n=3. Two-tailed unpaired t-test, **p<0.01. F, Relative FAM3C RNA expression level of 10-month WT and 5X FAD mice cortex. Data are presented as mean ± SD, n=3 mice. Two-tailed unpaired t-test, n.s., not significant. G, Relative FAM3C protein expression level of 10-month WT and 5X FAD mice cortex. Data are presented as mean ± SD, n=3 mice. Two-tailed unpaired t-test, *p<0.05. H, Relative FAM3C secretion level of iAst derived from healthy and AD patient iPSC. Data are presented as mean ± SD, n=3. Two-tailed unpaired t-test, ****p<0.0001. I-K, Exogenous FAM3C modulation on neuron growth, neurons were treated with FAM3C protein on DIV0, transfected with synapsin-GFP on DIV5, and measured on DIV7. I) representative confocal images of cortical neurons (DIV7) under different FAM3C treatments. J) sholl analysis of the neural complexity. K) the total number of intersections under different FAM3C treatments. Data are presented as mean ± SEM, n=30 neurons. Two-way ANOVA test (J) and one-way ANOVA test (K), **p<0.01, ****p<0.0001. L, Neuron viability after treatment with FAM3C on DIV8, DIV12, and DIV17 was measured by CellTiter-Glo Luminescent Assay. Data are presented as mean ± SD, n=3. Two-way ANOVA test, **p<0.01, ****p<0.0001.

We next determined the relative abundance of the 206 overlapped secreted proteins between TNF-α treated and vehicle-treated iAst. The secretion level of most overlapped proteins was downregulated by TNF-α treatment (Fig.5C). Among these proteins, we found that the secretion of FAM3 Metabolism Regulating Signaling Molecule C (FAM3C), a key molecule in epithelial-to-mesenchymal transition (EMT)^35^, was significantly down-regulated. qPCR and ELISA analysis further validated the decreased expression and secretion level of FAM3C in iAst upon TNF-α treatment (Fig.5D, E). Consistent with our observation, a recent study showed that the expression level of FAM3C was reduced in AD patients compared with age-matched controls^36^. FAM3C could bind to the γ-secretase complex to destabilize the β-secretase-cleaved APP carboxy-terminal fragment and reduce the secretion of Aβ40/42, indicating the potential clinical relevance of FAM3C in AD.

We further examined the expression level of FAM3C in 5X FAD mice and the AD patient iPSC-derived astrocytes. We found that the expression level of FAM3C was significantly decreased in 10-month-old but not in 5-month-old 5X FAD mice (Fig.5F, G, Fig.S8A, B), and the secretion level of FAM3C in AD-derived iAst was remarkably down-regulated as well (Fig.5H).

Previous studies have demonstrated that FAM3C is mainly expressed in neurons and might be protective in AD by decreasing Aβ40/42 secretion^35,36^. Our results showed that FAM3C was also expressed and secreted by astrocytes, and the secretion level of FAM3C was significantly decreased in AD-patient-derived iAst. We reasoned that the astrocyte secreted FAM3C might have a cell non-autonomous protection role on neurons. To test this hypothesis, we overexpressed the human APP in SH-SY5Y, treated it with recombinant human FAM3C protein, and measured Aβ40 and Aβ42 levels in the conditioned medium. However, the Aβ40/42 secretion was not changed (Fig.S8D), suggesting that the exogenous FAM3C may not participate in the metabolic regulation of Aβ in neurons.

To further investigate the function of astrocyte-secreted FAM3C, we incubated primary cultured cortical neurons with recombinant human FAM3C and evaluated the cell viability and the neurites’ complexity. We found that FAM3C could significantly enhance neurites’ complexity in a dose-dependent manner (Fig.5I-K). Moreover, FAM3C could also enhance neuronal viability (Fig.5L). Interestingly, this protective effect emerged only after two weeks of culture, indicating that FAM3C may play a crucial role in the long-term survival of mature neurons. Even though FAM3C had no significant influence on young neuron viability, it could promote neural development, as the complexity and numbers of total neurites and spines were increased after FAM3C treatment in DIV7 neurons (Fig.S8E). Our results demonstrated that astrocytes secreted FAM3C have a protective role on neurons, but it was downregulated in AD conditions.

### Secretome profiling of AD-patient (*ApoE 4/4*) derived astrocytes

Numerous studies have shown that astrocytes carrying neurodegenerative disorder-related mutations have non-cell autonomous toxicity on neurons^37–40^. As astrocytes are secretory cells, it has been proposed that this cell non-autonomous toxicity was mainly mediated through its secreted proteins. Indeed, we found that the viability of neurons incubated with AD-iPSCs derived astrocytes (AD-iAst, Fig.S9A) CM was dramatically decreased compared with that with healthy iAst CM (Fig.6A). Next, we profiled the secretome of the AD-iAst and identified 283 secreted proteins (Fig.S9B, Table S4). Pathway enrichment analysis revealed that AD-iAst secreted proteins are functionally enriched in neurological pathologies, particularly protein hemostasis (Fig.S9D), including serine/threonine kinase signaling pathway, response to TGF-β, APP metabolic process, receptor-mediated endocytosis, proteins processing, maintenance of protein location and response to external stimulus.

**Fig.6.**
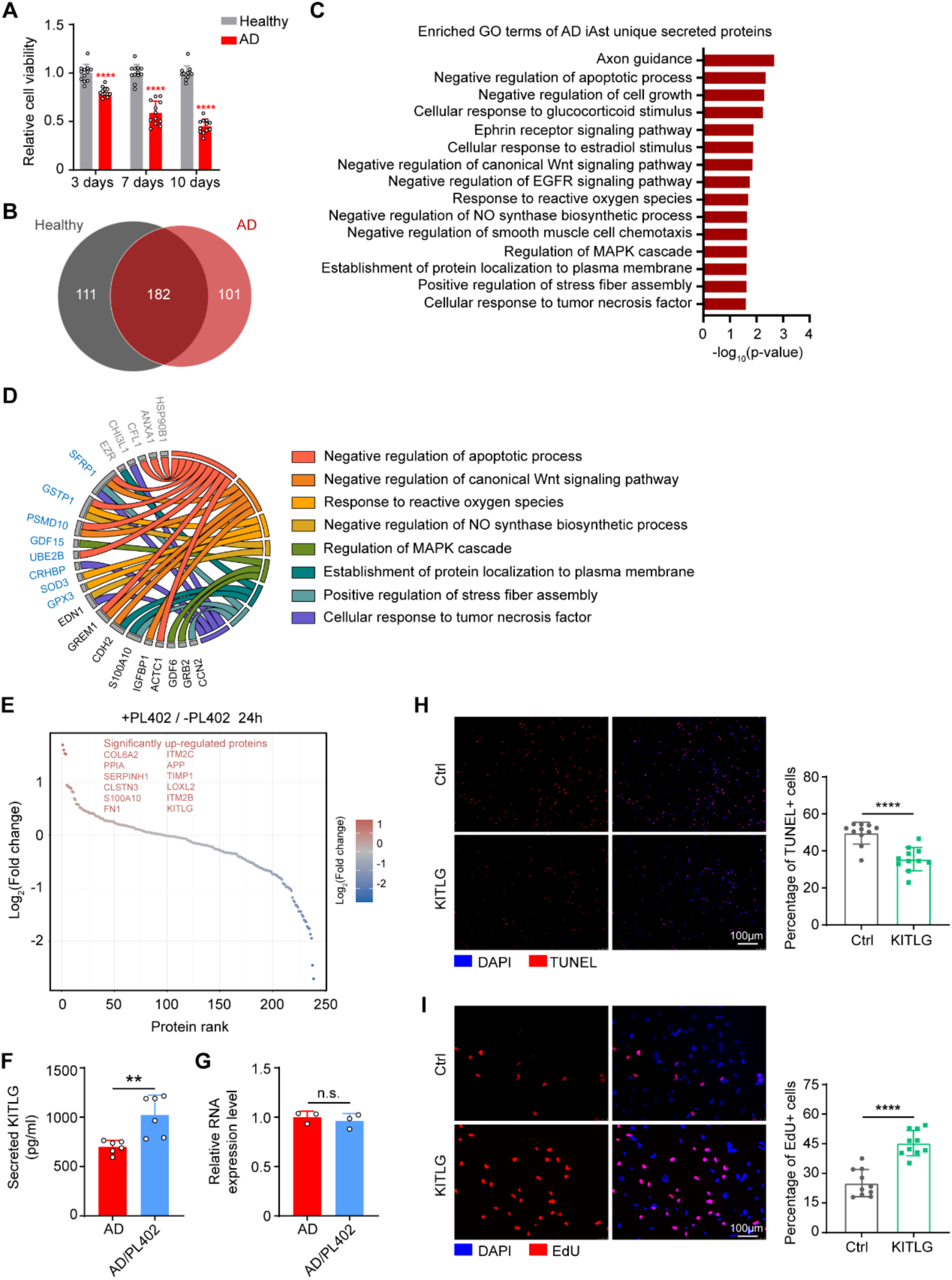
Secretome profiling of AD patient iPSC derived astrocytes. A, Effects of healthy and AD iAst conditioned medium on primary neuron cell viability. Data are presented as mean ± SD, n=3. Two-tailed unpaired t-test, ****p<0.0001. B, Comparison between AD iAst and healthy iAst secretome. AD iAst consists of 101 unique proteins and 182 overlapped proteins compared to the healthy iAst secretome. C, Top 16 enriched biological processes of 101 AD iAst unique secreted proteins according to DAVID database. D, Influence of PL402 treatment (300 mM) on the secretion of the 22 proteins enriched in highlight pathways. Gray, not detectable after PL402 treatment; blue, downregulated by PL402 treatment; black, not influenced by PL402 treatment. E, Relative quantitative analysis of the 182 overlapped secreted proteins between healthy and AD iAst secretome. F, Effects of PL402 treatment (300 mM) on AD iAst KITLG secretion. Data are presented as mean ± SD, n=3. Two-tailed unpaired t-test, **p<0.01. G, Relative AD iAst KITLG RNA expression level change after PL402 treatment. Data are presented as mean ± SD, n=3. Two-tailed unpaired t-test, n.s., not significant. H, Effects of KITLG treatment (10ng/ml) on neuronal survival. Neurons were treated with KITLG on DIV0 and measured on DIV15. Dead neurons were labeled with TUNEL (red). Data are presented as mean ± SD, n=11. Two-tailed unpaired t-test, ****p<0.0001. I, Effects of KITLG treatment (10ng/ml) on NPC proliferation. NPCs were treated with KITLG for four days; the proliferative cells were labeled with EdU(red). Data are presented as mean ± SD, n=11. Two-tailed unpaired t-test, ****p<0.0001.

The secretome between healthy iAst and AD-iAst differs a lot (Fig.6B). The AD-iAst specific secreted proteins were involved in biological processes that are implicated in the pathogenesis of AD, including negative regulation of the apoptotic process, response to reactive oxygen species, positive regulation of stress fiber assembly and cellular response to tumor necrosis factor, *etc* (Fig.6C), suggesting that the AD patient-derived astrocytes are indeed functionally impaired and pre-deposited for the development of AD pathology We next sought to determine whether our secretome profiling could be used to identify the potential mediator or molecular mechanism of therapeutic reagents. We treated the AD-iAst with PL402, a rhamnoside derivative recently shown to significantly attenuate Aβ pathology and cognitive defects in APP/PS1 transgenic mice^41^, for 24 hours and profiled the secretome after treatment (Table S5). We found that the secretion level of 13 out of 22 (59%) proteins related to AD pathogenesis are down-regulated (Fig.6D). Thus, the protective role of PL402 might be mediated by inhibiting the secretion of toxic proteins from mutant astrocytes to ameliorate cell non-autonomous toxicity.

We next quantitatively analyzed the relative change of all secreted proteins between PL402-treated or vehicle-treated AD-iAst and found that the secretion level of some proteins was significantly up-regulated (Fig.6E). We investigated the role of KITLG (SCF, stem cell factor) in the central nervous system. KITLG is a cytokine that binds to the c-KIT receptor and plays a critical role in hematopoiesis. We verified the up-regulated secretion of KITLG in PL402-treated AD-iAst with ELISA (Fig.6F). Interestingly, qPCR analysis of the expression level of KITLG demonstrated that the mRNA level of KITLG was not changed (Fig.6G), indicating that the up-regulation of KITLG protein secretion did not occur on the transcriptional level. It reinforced the necessity of studying secretomes under physiological and pathological conditions.

Next, we determined the function of KITLG on neurons *in vitro*. The primary cultured neurons were incubated with a medium supplemented with recombinant KITLG for 15 days, and the number of dying cells was evaluated by TUNEL staining. We found that KITLG treatment significantly improved cell viability, evidenced by reduced TUNEL-positive cells and increased survival cells in a Cell Viability Assay (Fig.6H, Fig.S10A, B). We further examined the effect of KITLG on the proliferation of neural progenitor cells (NPC) (Fig.S10C). Our results demonstrated that KITLG-treated NPCs exhibit a significantly increased EdU incorporation ratio than vehicle-treated NPCs (Fig.6I, Fig.S10D), suggesting that KITLG plays a vital role in NPC proliferation. Nevertheless, the function of KITLG and its contribution to the pathogenesis of neurological diseases such as AD merits further investigation.

## Discussion

Most drug candidates for neurological disorders succeed in mouse models but fail in human clinical trials. Thus, there is a strong need for human cellular models for disease mechanism studies and drug testing. Since astrocytes are crucial players in many neurological disorders and their secreted proteins are the primary mediator for the complex network of glia-neuron, glia-blood vessels, and glia-glia in health and disease, the ability to label and identify secretory proteins of human astrocytes will undoubtedly lead to new discoveries for the study of neurological disorders.

In this study, we have developed and optimized a method to label and identify secretory proteins from mAst and iAst. This method allowed us, for the first time, to directly profile and investigate the functions of secretory proteins of astrocytes on a large scale. Using this method, we found that in addition to the extracellular matrix organization, human astrocytes play critical roles in glycan metabolism, APP metabolic regulation and amyloid-beta formation, catabolic process, and clearance, *etc.* By comparing human iAst and mAst secretome profiling, we found that while astrocytes of both species share many secreted proteins and their relevant functions, each has its unique spectrum of properties. We further demonstrated the application of this method in disease mechanism studies and drug development by profiling the secretome of TNF-α treated iAst (to model neuro-inflammation) and a neuroprotective compound treated AD-iAst. Our method and the profiling of astrocyte secretome are valuable resources for studying the cellular mechanism of neurological diseases and the neuroscience community.

iPSC-derived astrocytes are the ideal tool for studying astrocyte biology and CNS drug development in a more disease-relevant setting. The previous serum-starvation-based culture methods for the secreted proteins collection and identification are problematic as the astrocytes are activated into a disease-like state in this culture condition. Therefore, it is difficult to identify authentic disease-induced changes. Our modified method and the iPSC-derived astrocyte secretome from healthy individuals and patients provide a basis for studying astrocyte mechanisms in disease versus health with the new expectations.

We compared the secretome of human iPSC-derived and primary cultured mouse astrocytes and found that many human astrocytes secreted proteins are not detected in mouse astrocytes. Even in the same cellular process, the proteins identified are different. From the perspective of proteomic analysis, it does not necessarily mean that mouse astrocytes do not secrete these proteins. However, some may provide insight into human astrocytes’ physiological and functional properties. Experimental approaches are undoubtedly needed for further investigation.

We identified over 101 differential secreted proteins from astrocytes derived from healthy individuals compared with those from AD (*APOE4/4*) patients derived iAst, and 43 from TNF-α treated iAst, which mimic the neuro-inflammatory conditions associated with neurological disorders such as AD and PD. A careful examination of the altered secretome in these diseases’ states may generate new insights into the role of astrocytes in the pathogenesis of neurological diseases, hoping to find new targets for diagnosis and prevention of the disease progression. We demonstrated this capability with FAM3C and KITLG, which we identified in secretome profiling of TNF-α treated healthy iAst and PL402-treated AD-iAst, respectively. FAM3C can be detected in the serum and cerebrospinal fluid and has been claimed to be a biomarker for AD^42^, whereas neuronal FAM3C plays a protective role for neurons in an AD mice model. Here, we show that astrocytes secreted FAM3C can promote neuronal survival and enhance neurite outgrowth, adding an additional layer of evidence for the potential clinical relevance of this protein with AD. The function of KITLG is well established in hematopoiesis, stem cell maintenance and gametogenesis, but its role in the adult brain is largely unknown. We show that the protein secretion, but not the mRNA level of KITLG, was up-regulated in the PL402-treated AD-iAst, and exogenous supplement of recombinant KITLG can significantly protect neurons from death and promote neural progenitor cell proliferation. In support of our *in vitro* data, the administration of PL402 can promote adult neurogenesis in an APP/PS1 AD mice model (Gang Pei et al., unpublished data).

As evidence accumulates for the importance of glia in the health and disease of CNS, the method for profiling and the datasets of the astrocytes secretome presented in this study will be a valuable resource for investigating the biology of human astrocytes and fruiting the development of novel diagnostic and treatment approaches for neurological disorders.

## Methods

*Methods and any associated references are available in the online version of the paper*.

*Note: Supplementary Information includes 8 figures, the list of the primers and antibodies used in this study is available online*.

## Ethics approval and consent to participate

*All housing, breeding, and procedures were approved by the Institutional Animal Care and Use Committees at the Interdisciplinary Research Center on Biology and Chemistry (IRCBC), Chinese Academy of Science*.

## Consent for publication

Not applicable.

## Availability of data and materials

The datasets during and/or analysed during the current study available from the corresponding author on reasonable request.

## Acknowledgements

We thank Prof. Gang Pei for kindly providing us with PL402 compound. We thank Prof. Yelin Chen for kindly providing us with APP695 plasmid. We thank all members of the Wang lab for helpful discussions.

## Author’s Contribution

J. L and J.G designed and performed the experiments and analyzed data; L. G helped with mass spectrum detection; G.M.M helped with bioinformatics analysis; all other authors provided technical support; W.Y.W, Y.Y.Z, J.C.H, and J.Z supervised the project.

## Materials and methods

### Cell culture and primary neuronal cell culture

HEK293 and C6 cells were maintained in high glucose DMEM media (Thermo Fisher Scientific, Cat# C11995500CP) supplemented with 10% heat inactivated FBS (Biological Industries, Cat# 04-001-1A) and 1% Pen/Strep (Thermo Fisher Scientific, Cat# 15140122) at 37°C and 5% CO_2_ atmosphere. Primary neuronal cells were cultured as previously described with certain modifications. Briefly, primary neurons from E16 ICR mice were plated on poly-L-lysine (Sigma, Cat# P6282) coated 24-well plate (for staining), 96-well plate (for cell viability assay) in plating media (Neurobasal, supplemented with 10% heat-inactivated FBS, 5mM Glutamax and 1% Pen/Strep) for 3h, neurons were maintained in regular media (Neurobasal (Thermo Fisher Scientific, Cat# 21103049), 1×B27 (Thermo Fisher Scientific, Cat# 17504044), supplemented with 5mM GlutaMax (Thermo Fisher Scientific, Cat# 35050061) and 1% Pen/Strep) at 37 °C and 5% CO_2_ atmosphere. Primary astrocytes from P1-3 ICR mice were plated on poly-D-lysine (Millipore, Cat# A-003E) coated T75 flask in high glucose DMEM media supplemented with 10% heat-inactivated FBS and 1% Pen/Strep at 37 °C and 5% CO_2_ atmosphere. The media was changed every 3 days. Astrocytes were shaken at 220 rpm for 12hr after reached confluency. Then the cells were plated for the follow-up experiments.

### Maintenance of human embryonic stem cell and induced pluripotent stem cells

Information of cell lines used in this study were summarized in Supplementary Table 8. Fibroblasts were reprogrammed with iPS 2.0 Sendai Reprogramming Kit (Thermo Fisher Scientific, Cat# A16517). iPSCs and ESC were incubated at 37^◦^C, 5% CO_2_, in matrigel (Corning, Cat# 354277) coated flasks, in StemFlex media (Thermo Fisher Scientific, Cat# A3349401), supplemented with 1% penicillin/streptomycin. Cells were passaged every 5 days with Gentle Cell Dissociation Reagent (STEMCELL, Cat# 07174), and assessed daily for confluence.

### Differentiation of human embryonic stem cell line into astrocytes

The process of differentiation followed a combination and adjustment of a former protocol^27^. Briefly, after reached confluence, ESC or iPSC were dissociated with 1mg/ml dispase II (Thermo Fisher Scientific, Cat# 17105041), seeded in untreated T25 flask to formed EBs. EBs were maintained in StemFlex media for the first three days. At day 4, change to neural differentiation media (NM) (DMEM/F12 (Thermo Fisher Scientific, Cat# 11330032), 1% Pen/Strep, 1% NEAA (Thermo Fisher Scientific, Cat# 11140050), 1X N2 (Thermo Fisher Scientific, Cat# 17502001), 20ug/ml heparin (Sigma, Cat# H3149)). At day 7, twenty to fifty intact EBs were seeded directly onto 20mg/ml laminin (Thermo Fisher Scientific, Cat# 23017015) pre-coated 6 well plates containing NM. On day 10, NM were added with morphogen RA (5 mM, Sigma, Cat# R2625), the media were changed every 3 days. At day 15, multiple rosette formations were present, and then were lifted and transferred to low attachment 6-well plates containing NM+RA to form astrospheres. On day 21, replaced medium with NM + EGF and FGF2 (20 ng/ml each, Peprotech, Cat# 100-47 and 100-18C) to expand the spheres. 75% of the medium were replaced every 2–3 days and spheres tightly attached to the well should be discarded. The spheres were maintained in this condition for up to 6months. For secretome profiling, spheres were dissociated into single cells with 0.05% Trypsin (Thermo Fisher Scientific, Cat# 25300062), and seeded in 10ug/ml PDL coated dishes and maintained in DMEM media supplemented with 10% FBS and 1% Pen/Strep until expanded to desired cell numbers.

### Differentiation of human embryonic stem cell line into neural progenitor cells (NPC)

Neural progenitor cell (NPC) was differentiated as previously described ^43^. In brief, hESC at 95–100% cell confluence was seeded onto Matrigel-coated 6-well plates in StemFlex media. The next day, medium was changed to N2B27 medium (50% DMEM/F12, 50% neurobasal, 0.5X N2, 0.5X B27, 1% Glutamax, 1% NEAA, 5 μg/mL insulin (Solarbio, Cat# I8830), and 1 μg/mL heparin) plus 5 μM SB431542 (Selleck, Cat# S1067) and 5 μM dorsomorphin (Selleck, Cat# S7840). The medium was changed every two days. After 8 days of induction, the cells were passaged on new six-well plates coated with Matrigel in N2B27 medium at a 1:2 ratio. The medium was changed every 2 days. After 16 days, canonical neural rosettes appeared. Then, these neural rosettes were dissociated to single cells with Accutase (STEMCELL, Cat# 07920). For EdU staining experiment, a total of 2.5 × 10^4^ NPCs was plated in a Matrigel-coated 24-well in N2B27 medium. The next day, the medium was changed to N2B27 medium, N2B27 medium supplemented with 10ng/ml KITLG (Abcam, Cat# Ab259391) or 300mM PL402 (a gift from Gang Pei Lab) respectively, the medium was changed every 2 days.

### Plasmid and lentivirus generation

Plasmid of hAPP is a gift from Yelin Chen lab. To generate lentivirus, expression vectors (pPGK HA-APP-FLAG) and packaging vectors (pMDL, pRev and pVSVG) were co-transfected into HEK293T cells at the ratio of 3:2:1:1. Medium containing lentivirus was collected 48 h later, sub-packaged and stored at -80 °C until use.

### Immunofluorescence staining

Cells were fixed in 4% PFA (Solarbio, Cat# P1110) for 30 min at room temperature and blocked in blocking media (5% normal goat serum, 0.2% Triton X-100 in PBS) for 1h at room temperature. Cells were incubated with primary and secondary antibodies diluted in antibody dilution media (1% normal goat serum, 0.1% Triton X-100 in PBS) overnight at 4°C and 1h at room temperature, respectively, followed by washing four times with PBS. Coverslips were mounted onto glass slides with ProLong™ Gold Antifade Mountant medium containing DAPI (Thermo fisher scientific, Cat# P36931). Images were taken with Leica SP8 confocal microscope (Leica Co., Germany). Neuronal complexity was measured with Sholl Analysis (Fiji).

### Western blotting

For brain sample, mice brain dissected was harvested and lysed in RIPA buffer (Beyotime, Cat# P0013B). For cell sample, cell was harvested and lysed in 8M urea/100mM Tris buffer. Then, the sample was sonication followed by centrifuging for 10 min at 20,000g at 4°C and the supernatant was collected for SDS-PAGE. Membrane was blocked with 5% (m/v) nonfat milk for 1h at room temperature followed by immunoblotting with indicated primary and secondary antibodies overnight at 4°C and 1h at room temperature, respectively. Antibodies are listed in Supplementary Table 7. For silver staining, the experiment was conducted follow the manufacturer’s (Beyotime, Cat# P0017S) instructions.

### Pulse labeling of cells with AHA and two-step peptide level enrichment

When isolating secreted proteins in serum-containing media, we found that high serum concentration (10%) can dramatically impact the efficiency of click reaction (data not shown). Therefore, we optimized the culture conditions to ensure that neither the click reaction efficiency nor the cell growth was affected. Our results show that 24h of 0.5% serum culture is the most suitable condition (data not shown). If not mentioned, the follow-up experiments were all conducted under 0.5% serum.

To deplete cells of methionine, lysine and arginine, the cells were washed 3 times with warmed PBS. Then, the cells were labeled with AHA in DMEM without methionine, arginine and lysine (Thermo fisher scientific, Cat# 21013024) with 0.5% dialyzed FBS, 1% Glutamax, 100uM l-cystine and 1% Pen/Strep supplemented with 50 uM l-AHA (AnaSpec, Cat# AS-63669) for 24 hours. Collected media was then concentrated with Amicon Ultra Centrifugal Filters 3-kDa cutoff (Millipore, Cat# UFC900324), mixed with 8M urea/100mM Tris (1:1) and EDTA free protease inhibitor (Roche, Cat# 11873580001). Click reaction were conducted according to a former study^44^ with certain adjustment. For each 1 ml of the supernatant, add 2 μl triazole ligand (200 mM, prepared as described previously^45^), 4 μl biotin-PEG4-alkyne (25mM, Sigma, Cat# 764213) and 20 μl of CuBr suspension (10μg/ml, Sigma, Cat# 254185). Incubated samples for 18 h at 4°C with constant agitation. We then pull down these biotin-labeled proteins using Pierce Monomeric Avidin Agarose (Thermo fisher scientific, Cat# 20228) overnight at 4°C with constant agitation. Instead of directly on-beads digestion, we next eluted these resin binding proteins with 2% trifluoroacetic acid (TFA, Sigma, Cat# 6508), 50% acetonitrile (ACN, Sigma, Cat# 900667).

The eluted proteins were digested into peptides with Trypsin (Promega, Cat# V5111), and underwent the second step peptide level enrichment. At last, we eluted these biotin-labeled peptides with 2% TFA, 50% ACN. The eluted fraction was dried by centrifugation under a vacuum at 4°C and restored at -80°C then resuspended in 0.1% FA (v/v) for the subsequent LC-MS/MS analysis.

### LC-MS/MS analysis

The peptide mixture was analyzed using an on-line EASY-nL-LC 1000 coupled with an Orbitrap Q-Exactive HF mass spectrometer (Thermo Scientific, USA). The sample was loaded directly onto a 15-cm home-made capillary column (C18-AQ, 1.9 um, Dr.Maisch, 100 μm I.D.). Mobile phase A consisted of 0.1% FA, 2% ACN and 98% H_2_O and mobile phase B consisted of 0.1% FA, 2% H2O and 98% ACN. A 60 min gradient (mobile phase B: 4 % at 0 min, 8 % at 4 min, 22 % at 45 min, 30 % at 53 min, 90 % at 57 min, 90 % at 60 min) was used at a static flow rate of 300 nl/min. The data were acquired in a data-dependent (top-20) mode. High-energy collisional dissociation (HCD) was used to fragment the precursor peptides. For MS1, the scan range was set to 350 – 1,500 m/z at a resolution of 60,000. The AGC target was set as 3e6 with a maximum injection time of 20 ms. For MS2, the resolution was set to 30,000 with a fixed first mass of 120 m/z. The AGC target was set to 1e5 and the maximum injection time was set to 100 ms. Each sample was analyzed twice.

### MS Data Analysis

Maxquant software (version 1.5.2.8) was used for quantifications. The MS data were searched against the SwissProt human protein database (downloaded August 2018, 20,191 entries) and the build-in contaminant protein list. Trypsin was set as the enzyme, and the maximum missed cleavage was set to 2. Carbamidomethyl (C) (+57.02 Da) was set as a fixed modification and oxidation of methionine (+15.99 Da), acetylation of protein N termini (+42.01 Da) were set as variable modifications. In addition, we substitute AHA (−4.98 Da) and biotin-AHA (+452.23 Da) for methionine. “Match between runs” was applied. A false discovery rate of 1% was set at protein, peptide levels.

### ELISA

The supernatants collected were assayed by Human FAM3C ELISA Kit (FInetest, Cat# EH8412), Human KIT ligand ELISA Kit (Jonln, Cat# JL19115), Amyloid beta 40 Human ELISA Kit (Thermo Fisher Scientific, Cat# KHB3481) and Amyloid beta 42 Human ELISA Kit (Thermo Fisher Scientific, Cat# KHB3441) according to the manufacturer’s instructions respectively.

### RNA isolation and qPCR

Total RNA was extracted with TRIzol reagent (Thermo fisher scientific, Cat# 15596018) following manufacturer’s instructions and reverse transcribed with Hifair® 1st Strand cDNA Synthesis SuperMix for qPCR (gDNA digester plus) (Yeason, Cat# 11121ES60). cDNA was quantified using TB Green® Premix Ex Taq™ (Tli RNase H Plus) (Takara, Cat# RR420B) with an Applied Biosystems™ QuantStudio™ 6 Flex Real-Time PCR System (Thermo fisher scientific, USA) following manufacturer’s instructions.

### EdU staining

An EdU assay were performed with Click-iT™ Plus EdU Cell Proliferation Kit for Imaging, Alexa Fluor™ 594 dye (Thermo fisher scientific, Cat# C10339) according to the manufacturer’s instructions. Astrocytes were treated with medium containing EdU (10μM) for 4h, and NPCs were treated with EdU (10μM) for 16h.

### Cell viability measurement

Cell viability was detected with CellTiter-Lumi Luminescent Cell Viability Assay Kit (Beyotime, Cat# C0065M) and Cell Counting Kit-8 kit (Beyotime, Cat# C0037) according to the manufacturer’s instructions.

### Function enrichment analysis

GO Function enrichment analysis was performed using DAVID 6.8 web tools (https://david.ncifcrf.gov). Pathway enrichment analysis was performed using g:Profiler web server (http://biit.cs.ut.ee/gprofiler/gost) according to a previously published protocal^46^.

### Statistical analysis

The statistical analysis was performed using GraphPad Prism 8 software. All results were expressed as mean ± SD with P <0.05 indicating significance.

**Supplementary Table 6.**
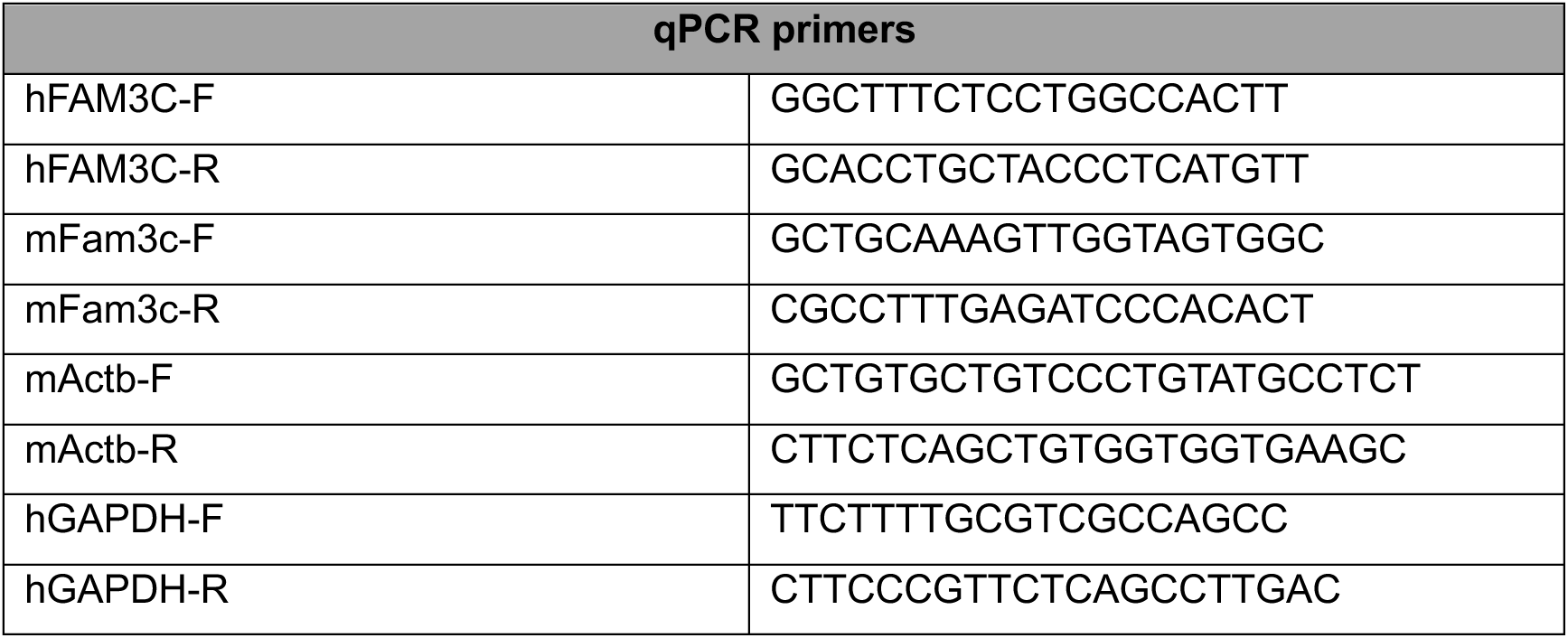
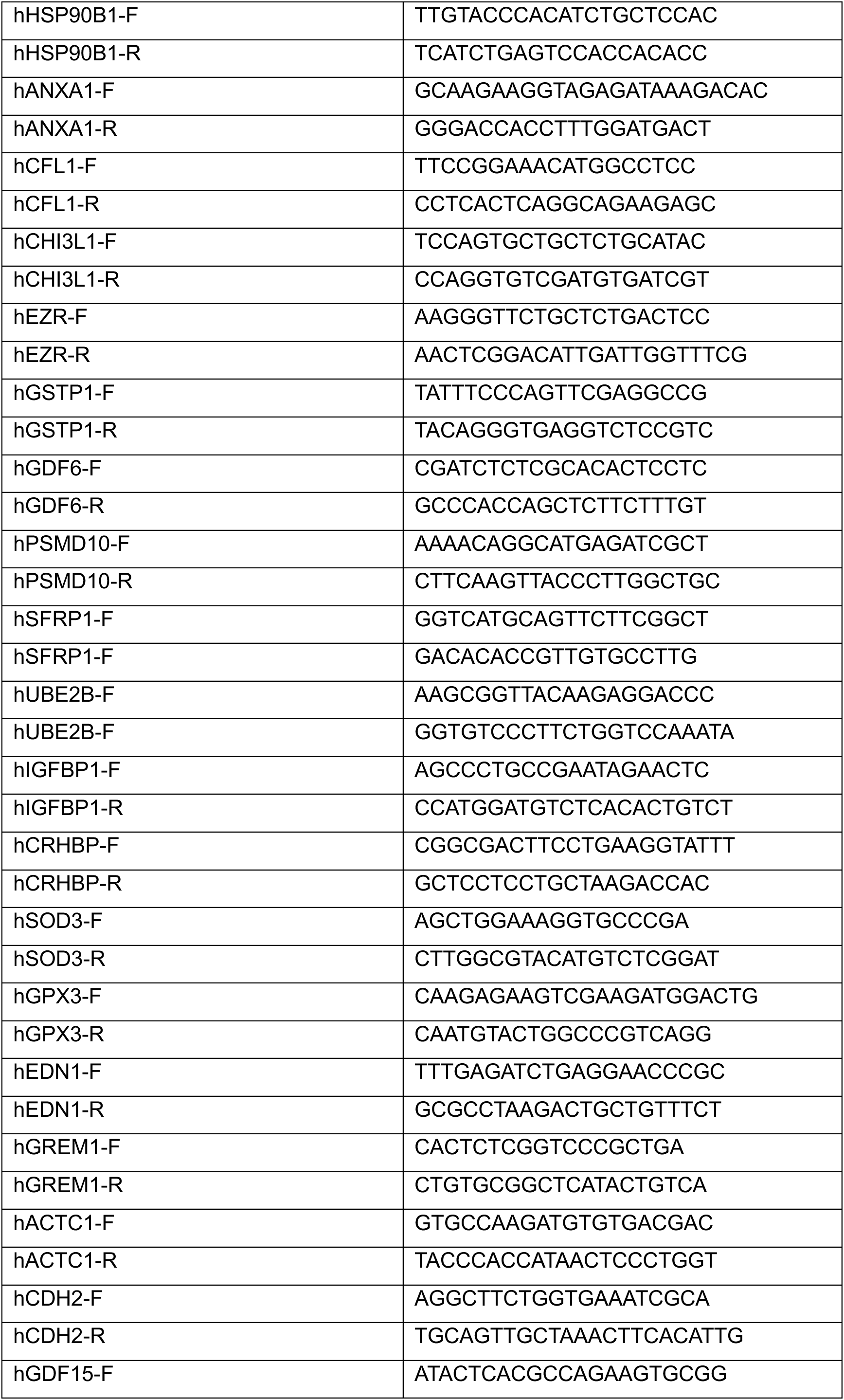
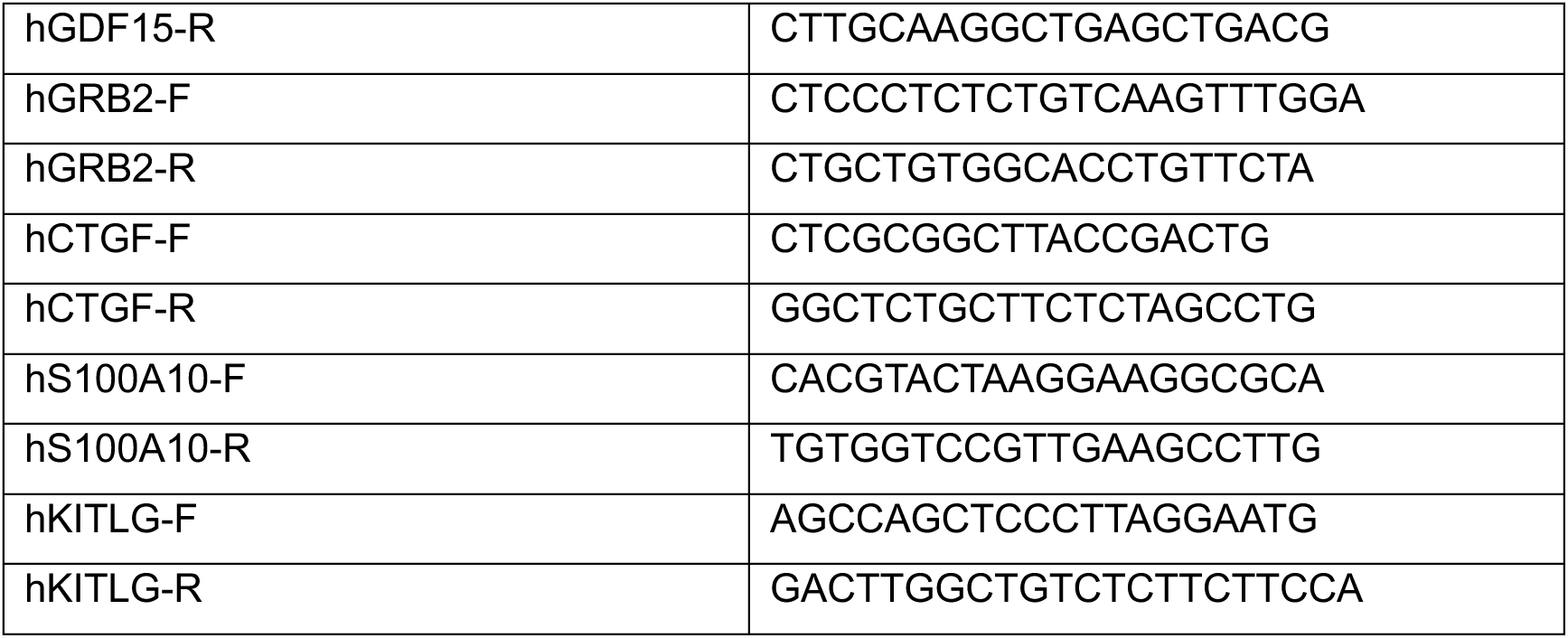
qPCR primers used in the study

**Supplementary Table 7.**
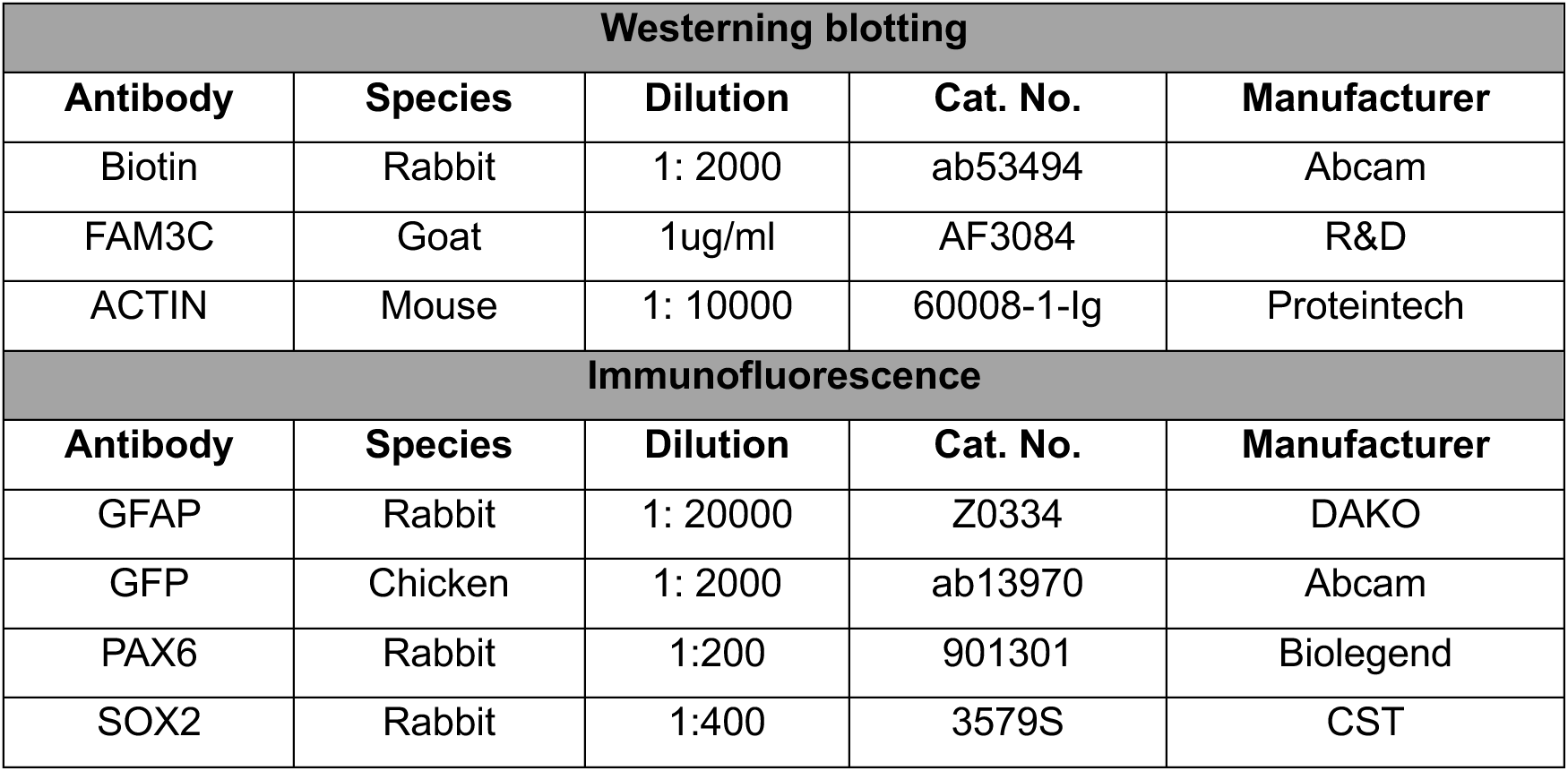
Antibodys used in the study

| Westerning blotting |  |  |  |  |
| --- | --- | --- | --- | --- |
| Antibody | Species | Dilution | Cat. No. | Manufacturer |
| Biotin | Rabbit | 1: 2000 | ab53494 | Abcam |
| FAM3C | Goat | 1ug/ml | AF3084 | R&D |
| ACTIN | Mouse | 1: 10000 | 60008-1-Ig | Proteintech |
| Immunofluorescence |  |  |  |  |
| Antibody | Species | Dilution | Cat. No. | Manufacturer |
| GFAP | Rabbit | 1: 20000 | Z0334 | DAKO |
| GFP | Chicken | 1: 2000 | ab13970 | Abcam |
| PAX6 | Rabbit | 1:200 | 901301 | Biolegend |
| SOX2 | Rabbit | 1:400 | 3579S | CST |

**Supplementary Table 8.** Cell line used in this study

| ESC and iPSC line |  |  |  |  |
| --- | --- | --- | --- | --- |
| Cell ID | Gender | Age of collection | Mutation | Disease relevance |
| H1 | M | Blastocyst stage | APOE3/3 | Healthy |
| ZYF | F | 74 | APOE 4/4 | AD |

## Supplementary information

**Fig.S1.**
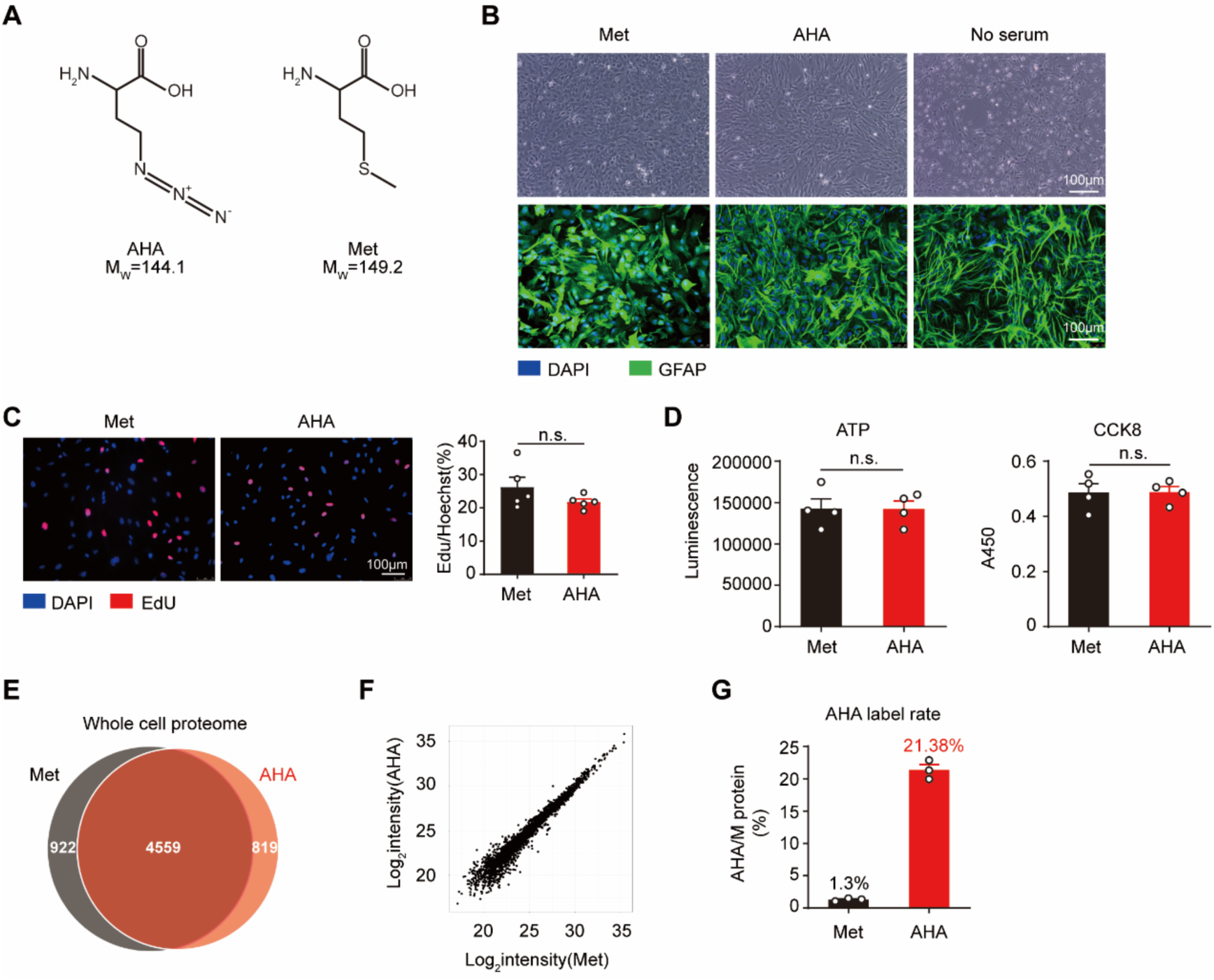
Influence of AHA replacement culture on astrocytes statuses. A, Structure and molecular weight of AHA and Met. B, Representative images of astrocytes after no serum or AHA treatment, astrocytes were labeled with GFAP (green) antibody. C, Effects of AHA treatment on astrocytes proliferation rate after 24h treatment, proliferative cells were labeled with EdU (red). Data are presented as mean ± SD, n=5. Two-tailed unpaired t-test, n.s., not significant. D, AHA has no significant influence on cell viability after 24h treatment. Data are presented as mean ± SD, n=4. Two-tailed unpaired t-test, n.s., not significant. E,F, AHA treatment has no significant influence on whole cell proteome after 24h treatment, correlation coefficient =0.9774626. G, 21.38% of the protein can be labeled by AHA after 24h treatment. Data are presented as mean ± SD, n=3.

**Fig.S2.**
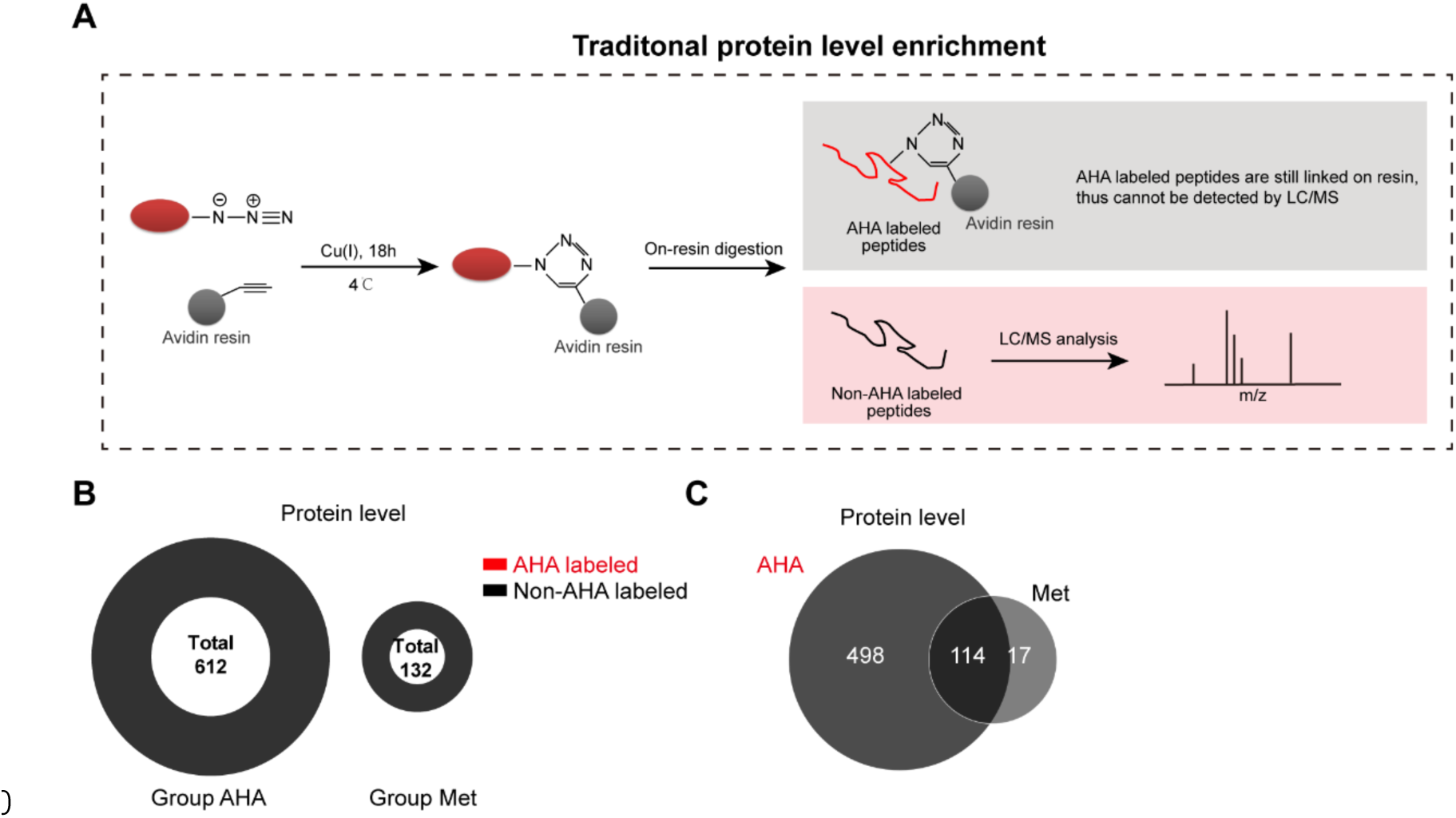
Schema of protein level enrichment. A, Schema of protein level enrichment. AHA labeled proteins were selectively and covalently linked to alkyne-activated resin via click chemistry reaction. Then the isolated proteins were digested on beads for the following mass spectrometry analysis. As AHA labeled peptides are still linked on resin, only the non-AHA labeld peptides can be detected during the subsequent LC/MS analysis. B, Numbers of proteins profiled by protein level enrichment. No AHA labeled proteins were detected in both groups. C, Comparison of the profiled proteins between the AHA and Met group.

**Fig.S3.**
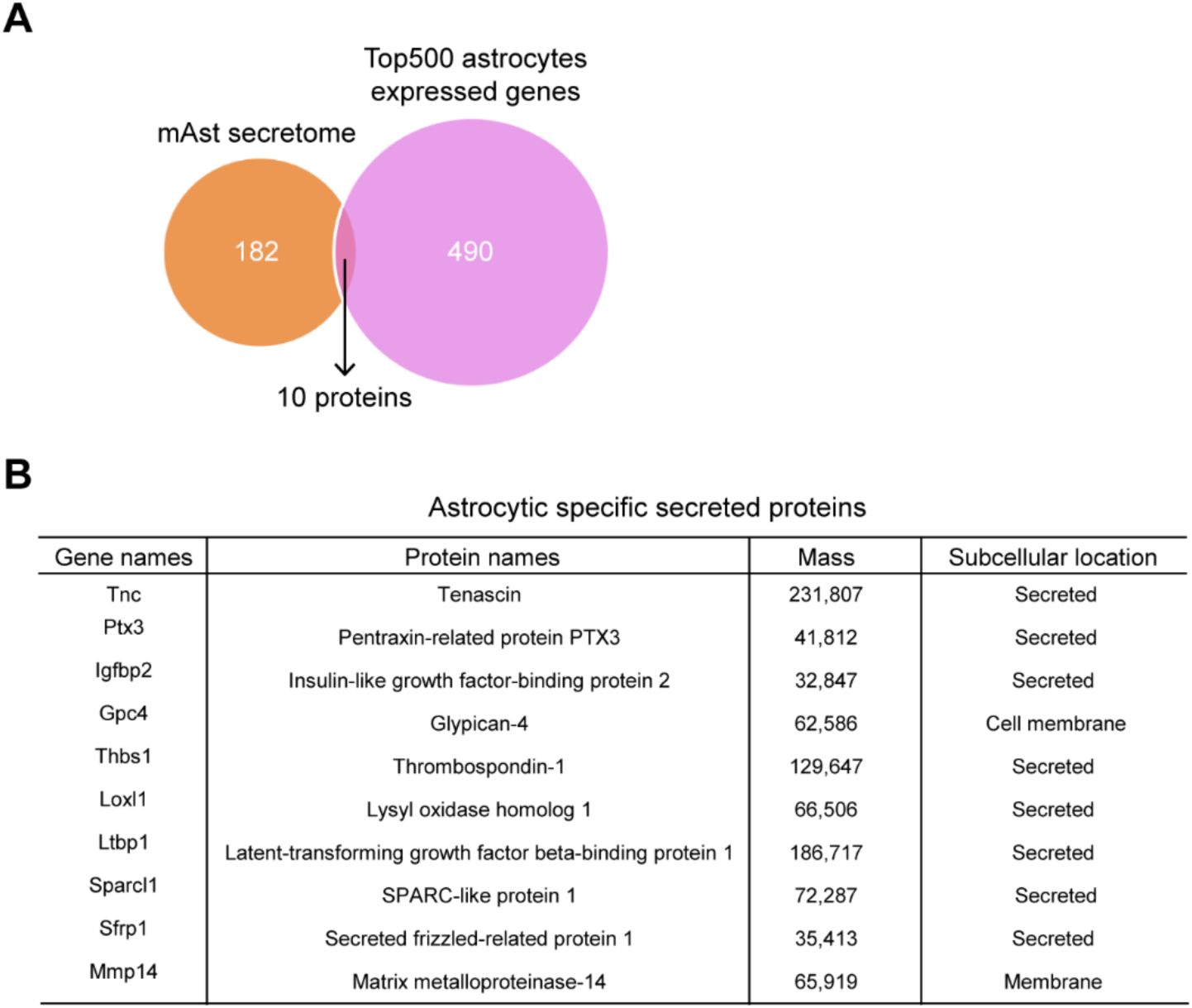
Comparison between mAst secretome and top500 astrocytes expressed genes. a, Venn plot of the mAst secretome and top500 astrocytes expressed genes. b, The detailed information of the 10 overlapped proteins.

**Fig.S4.**
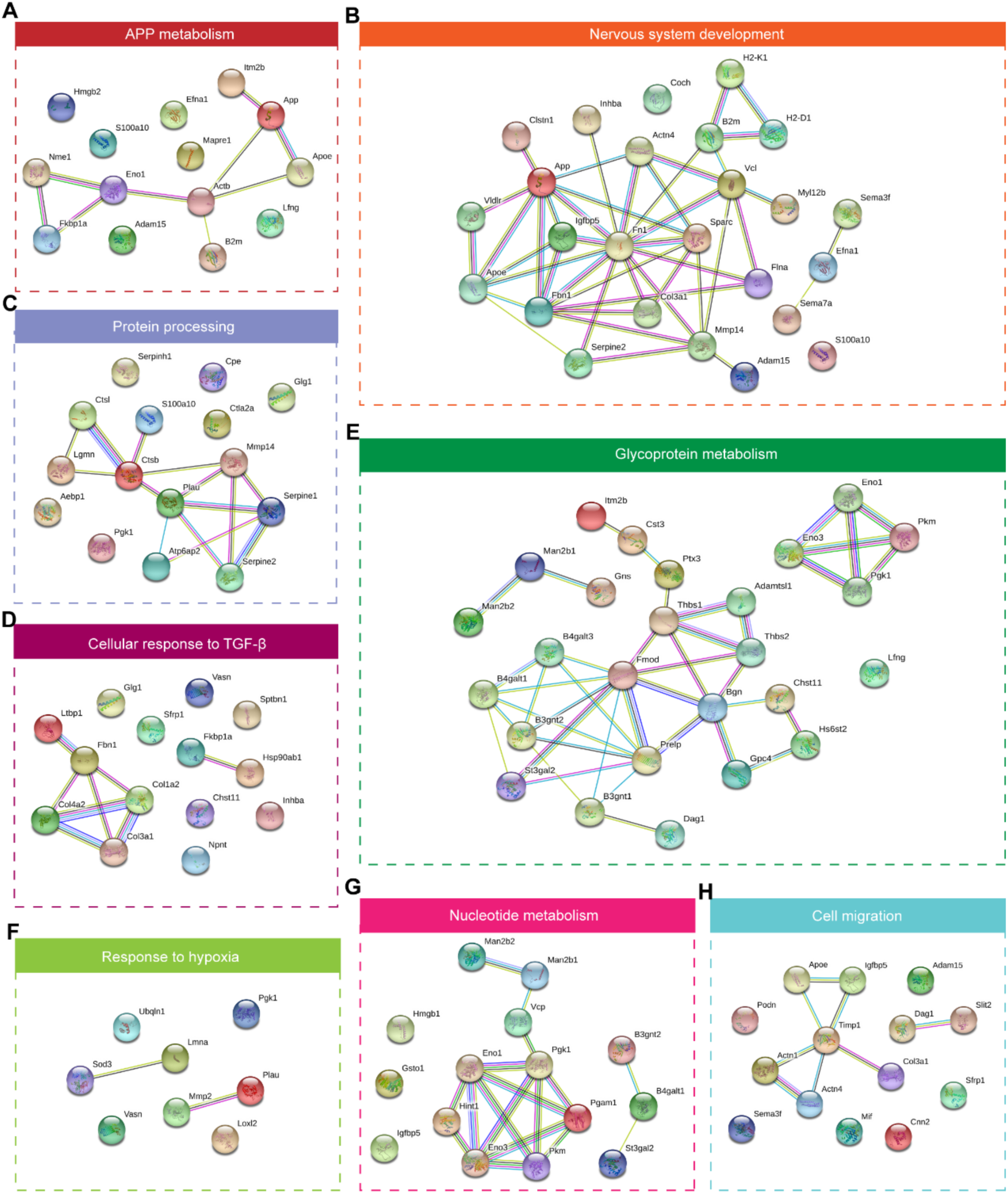
Protein-protein interaction analysis of mAst secretome. Protein-protein interaction analysis of mAst secreted proteins enriched in APP metabolism(A), nervous system development(B), protein processing(C), cellular response to TFG-β(D), glycoprotein metabolism(E), response to hypoxia(F), nucleotide metabolism(G) and cell migration (H) pathways.

**Fig.S5.**
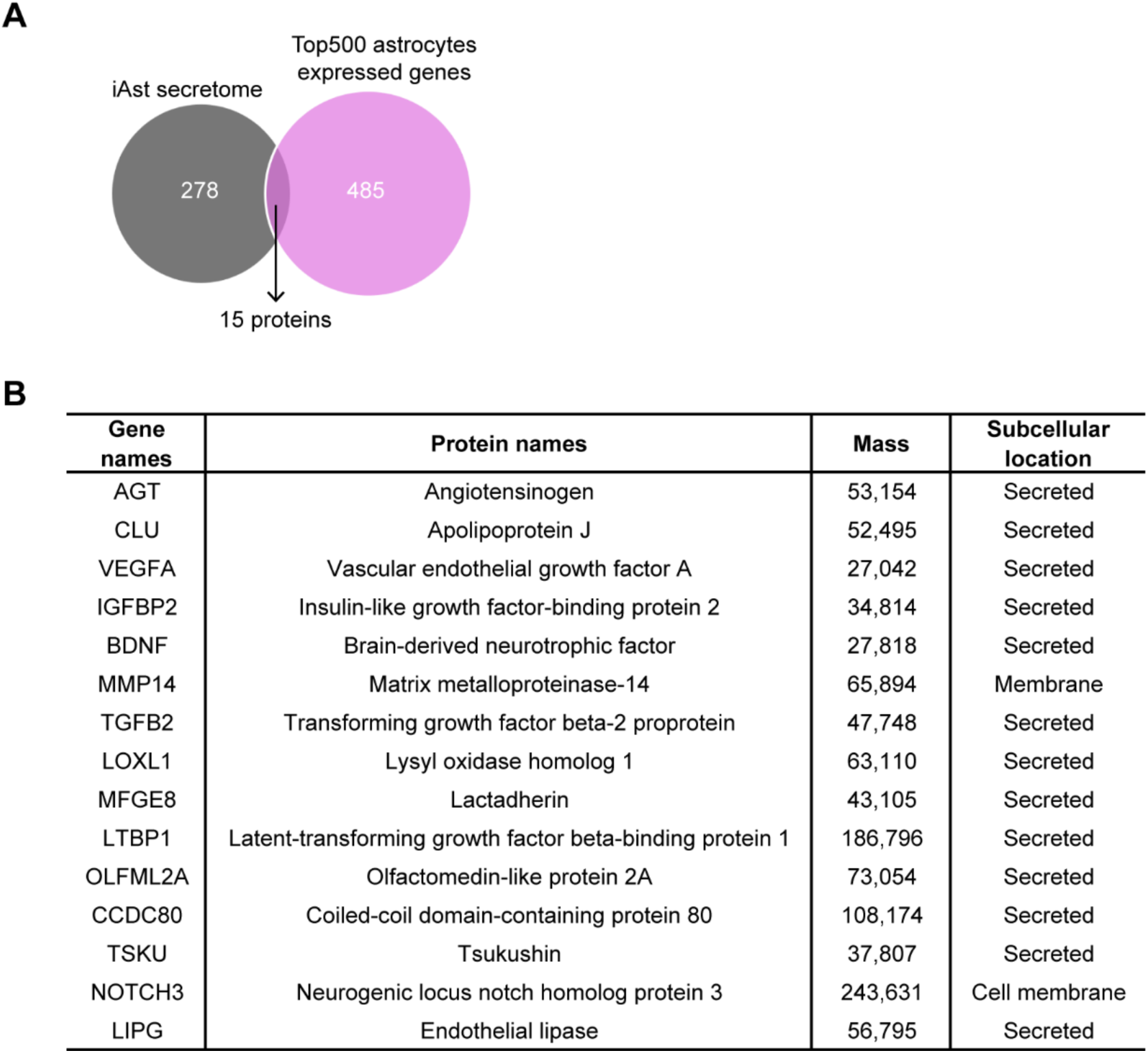
Comparison between iAst secretome and top500 astrocytes expressed genes. A, Venn plot of the iAst secretome and top500 astrocytes expressed genes. B, The detailed information of the 15 overlapped proteins.

**Fig.S6.**
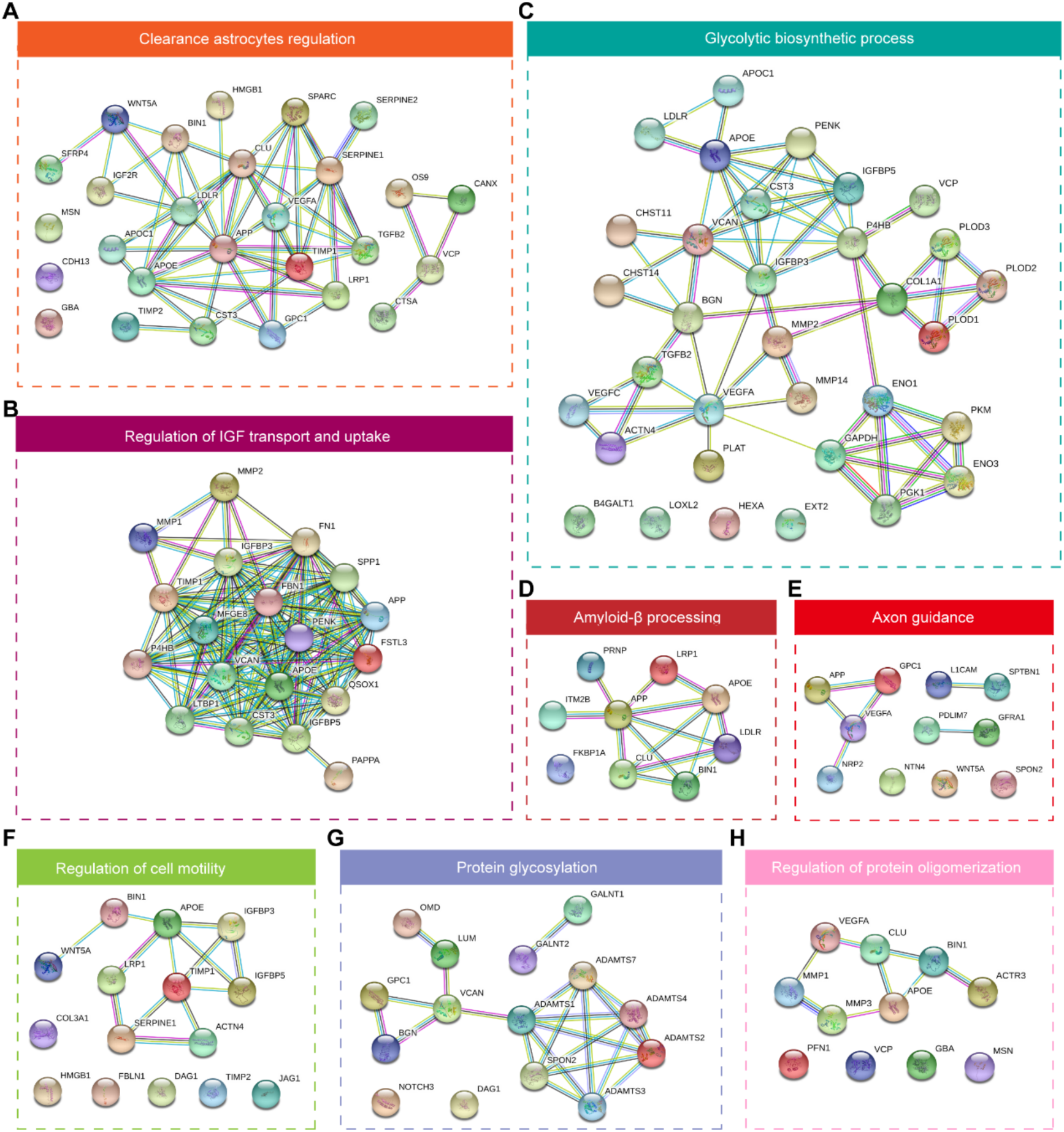
Protein-protein interaction analysis of the iAst secretome. Protein-protein interaction analysis of the iAst secreted proteins enriched in clearance astrocytes regulation(A), regulation of IGF transport and uptake(B), glycolytic biosynthetic process(C), amyloid-β processing(D), axon guidance(E), regulation of cell motility(F), protein glycosylation(G) and regulation of protein oligomerization(H) pathways.

**Fig.S7.**
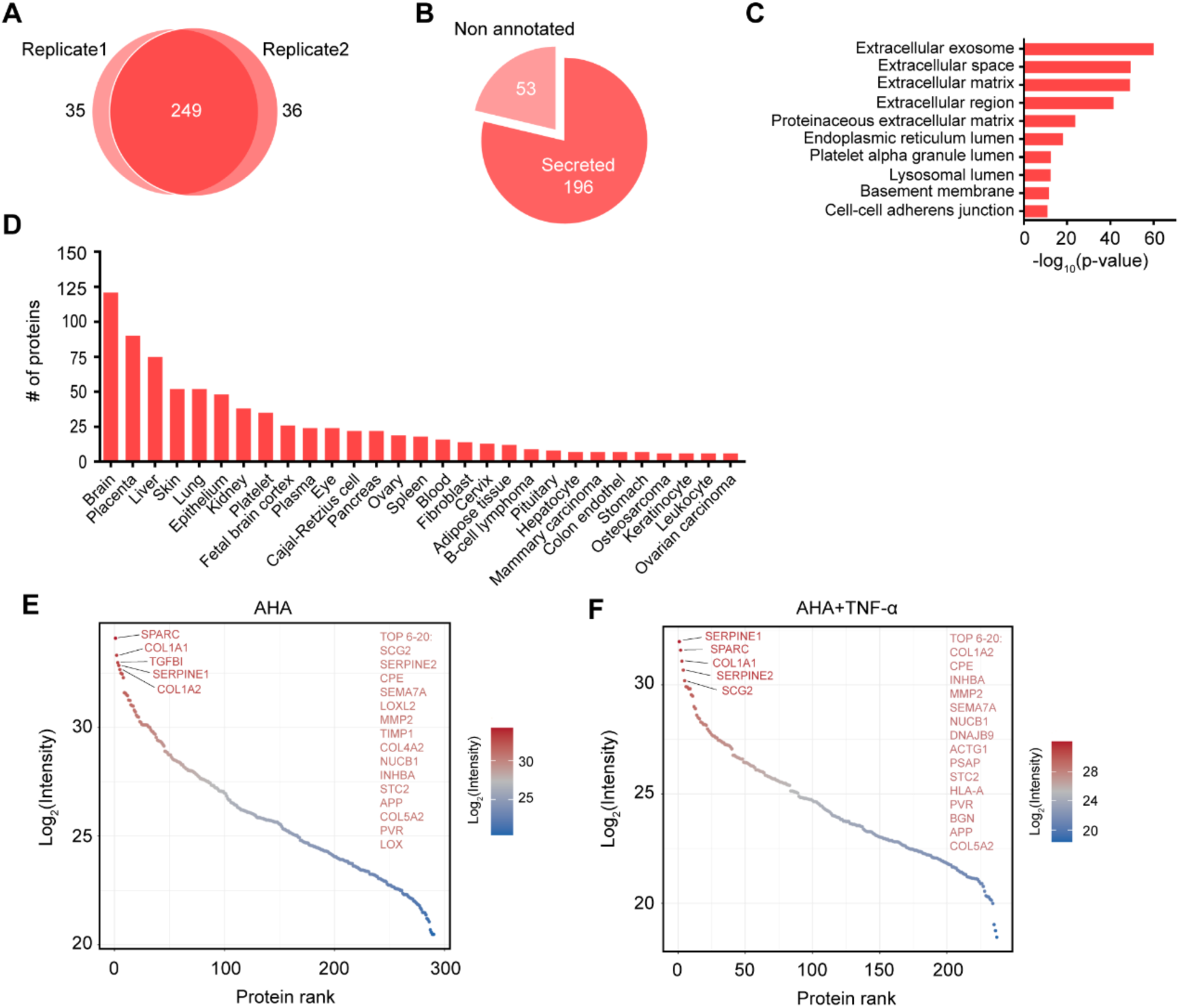
iAst secretome profiling after TNF-α treatment. A, TNF-α treated iAst secretome was also analyzed in duplicate by LC-MS/MS. Greater than 87% of the qualified proteins were found in both replicates. N=2 biological replicates. B, Statistical analysis of the subcellular location of the proteins detected in both replicates. Proteins were annotated as secreted, extracellular space and membrane in Uniprot database were all regard as secreted in our statistics. C, Cellular component distribution of the identified secretome proteins was analyzed by DAVID database, almost all the top 10 enriched terms were related to extracellular location. D, Tissue distribution of TNF-α treated iAst secretome according to DAVID database. E,F, Ranked intensity of TNF-α untreated(E) and treated(F) iAst secretome. Top20 secreted proteins of each group were annotated.

**Fig.S8.**
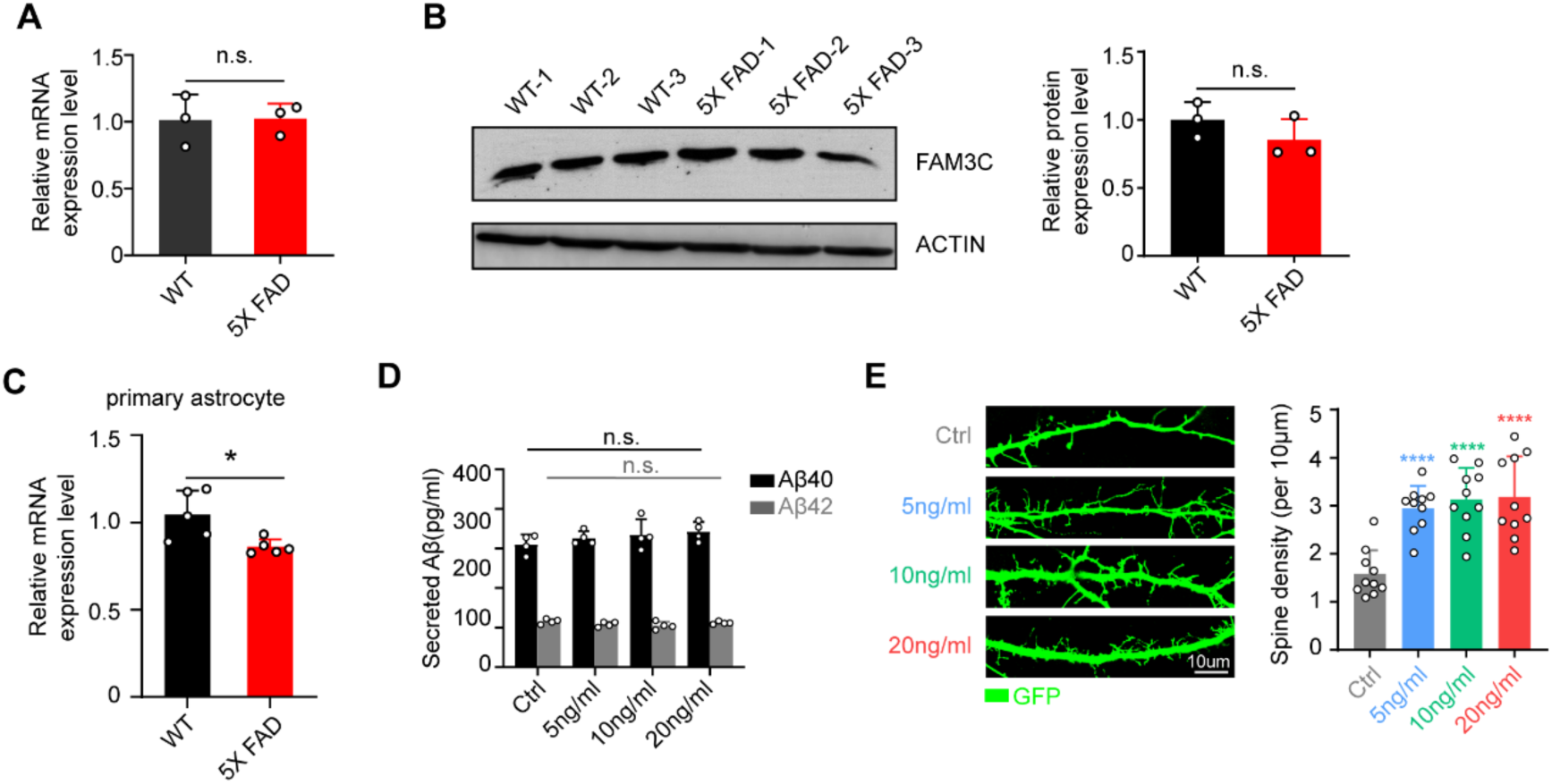
FAM3C expression in AD model, related to Fig.5. A, Relative FAM3C RNA expression level in the cortex of 5-month WT and 5X FAD mice. Data are presented as mean ± SD, n=3 mice. Two-tailed unpaired t-test, n.s., not significant. B, Relative FAM3C protein expression level in the cortex of 5-month WT and 5X FAD mice. Data are presented as mean ± SD, n=3 mice. Two-tailed unpaired t-test, n.s., not significant. C, Relative FAM3C RNA expression level of WT and 5X FAD mice primary astrocytes. Data are presented as mean ± SD, n=3. Two-tailed unpaired t-test, *p<0.05. E, FAM3C modulation of Aβ40/42 secretion in SH-SY5Y cell line which overexpressed hAPP. Data are presented as mean ± SD, n=3 mice. Two-tailed unpaired t-test, n.s., not significant. E, FAM3C modulation on neuronal spine density, neurons were treat with FAM3C protein on DIV0, transfected with synapsin-GFP on DIV5 and measured on DIV7. Data are presented as mean ± SD, n=10. One-way ANOVA test, ****p<0.0001.

**Fig.S9.**
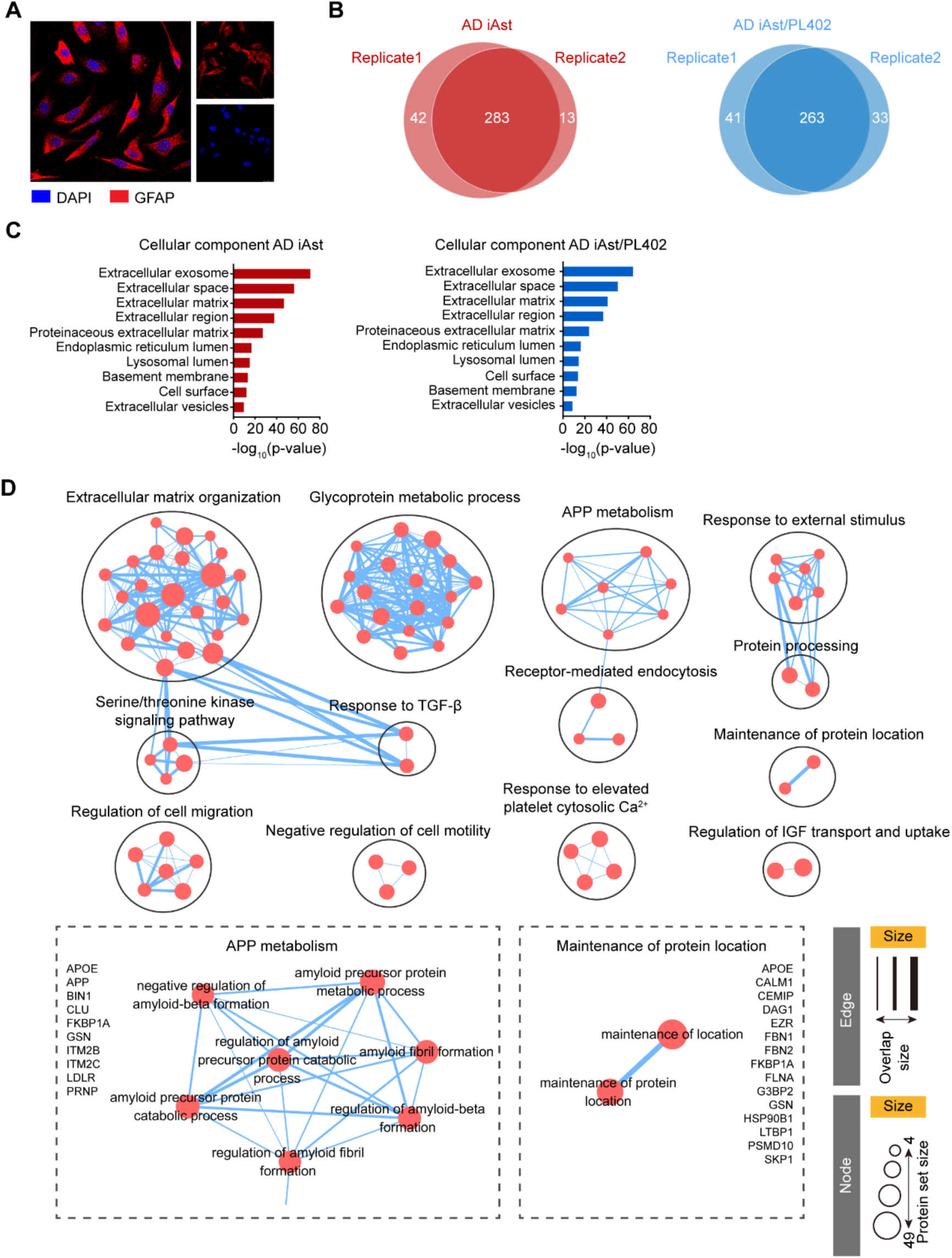
Secretome profiling of AD-iAst. A, Typical confocal image of induced astrocytes derived from AD patient iPSC, astrocytes were labeled with GFAP (red). B, AD iAst secretome was also analyzed in duplicate by LC-MS/MS. Greater than 90% of the qualified proteins were found in both replicates. N=2 biological replicates. C, Cellular component distribution AD iAst secretome proteins according to DAVID database, top 10 enriched terms were listed here. D, Pathway enrichment analysis of AD iAst secretome.

**Fig.S10.**
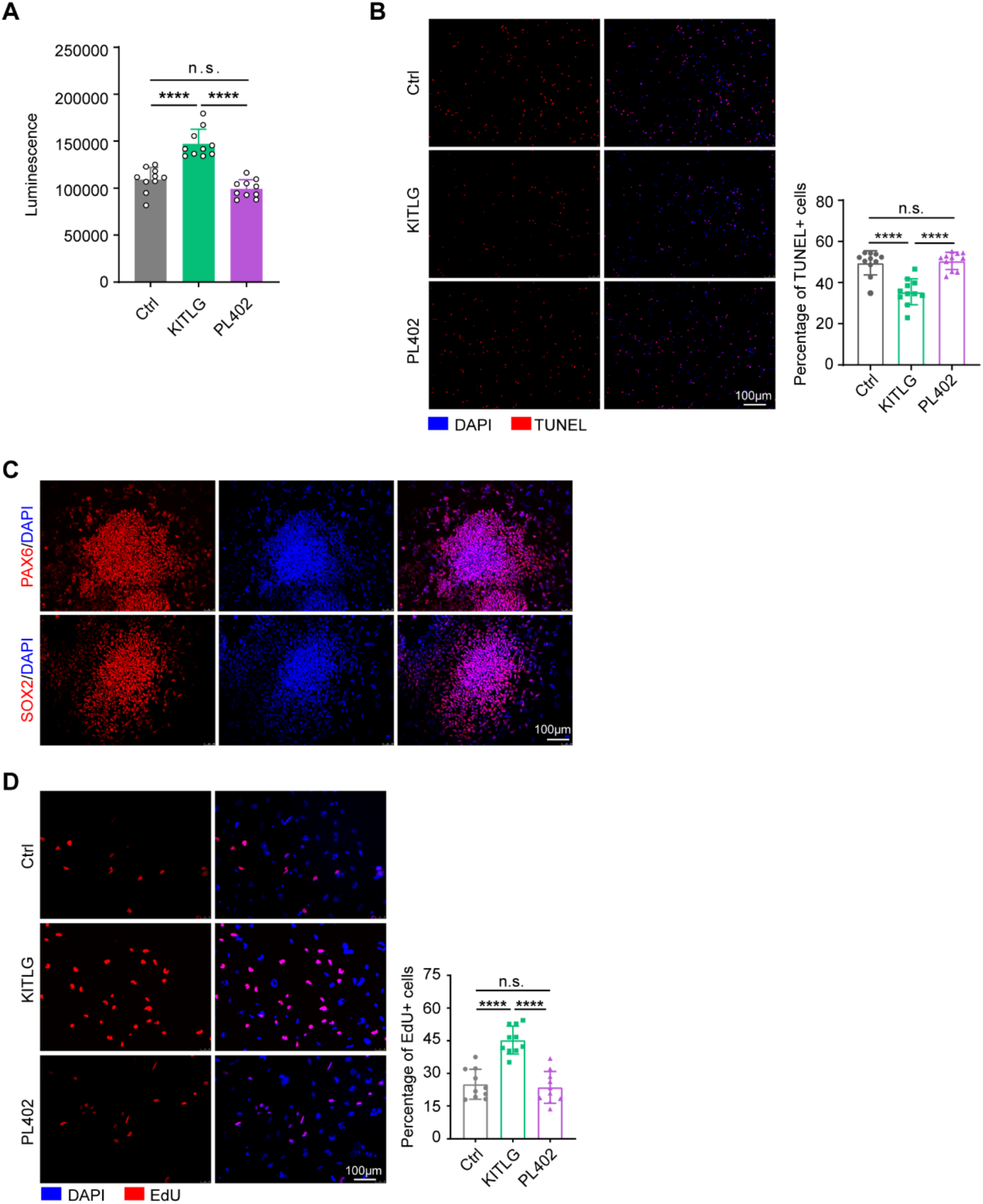
The neurotrophic effects of KITLG. A, Influence of KITLG (10ng/ml) and PL402 (300mM) on neuronal cell viability on DIV15. Data are presented as mean ± SD, n=3. One-way ANOVA test, ****p<0.0001. B, Influence of KITLG (10ng/ml) and PL402 (300mM) treatment on neuronal survival on DIV15, dead neurons were labeled with TUNEL (red). Data are presented as mean ± SD, n=11. One-way ANOVA test, ****p<0.0001, n.s., not significant. C, Typical confocal images of NPC. NPC were labeled with PAX6 (red) and SOX2 (red). D, Influence of KITLG (10ng/ml) and PL402 (300mM) treatment on NPC proliferation, proliferative cells were labeled with EdU (red). Data are presented as mean ± SD, n=11. One-way ANOVA test, ****p<0.0001, n.s., not significant.

